# HeteroRC: Decoding latent information from dynamic neural responses with interpretable heterogeneous reservoir computing

**DOI:** 10.64898/2026.04.04.716475

**Authors:** Runhao Lu, Sichao Liu, Yanan Liu, John Duncan, Richard N. Henson, Alexandra Woolgar

## Abstract

Time-resolved neural decoding is widely used to track information represented in neural activity, but conventional linear decoders primarily capture phase-locked evoked responses and often fail to recover representations embedded in nonlinear or non–phase-locked dynamics, potentially limiting the interpretation of neural coding. Here, we introduce HeteroRC, a biologically inspired and interpretable decoding framework based on heterogeneous reservoir computing. HeteroRC projects neural signals into a high-dimensional recurrent state space with heterogeneous time constants, enabling nonlinear feature expansion and multiscale temporal integration directly from raw neural time series. Simulations demonstrate that HeteroRC significantly outperforms linear decoders and a suite of artificial neural networks (including RNNs, LSTMs, Transformers and EEGNet) on evoked responses while robustly capturing induced oscillatory power, phase synchrony, and aperiodic modulations—dynamics that are largely latent to conventional linear methods. We further validate HeteroRC on two empirical EEG datasets. In a motor imagery task, it substantially improves decoding accuracy and exhibits superior cross-temporal generalisation, revealing dynamic representational transformations. In an attentional priority task, HeteroRC uncovers statistically learned spatial priority information that remains hidden from conventional methods, successfully decoding these latent states previously thought to be ‘activity-silent’. Furthermore, we develop a dual-level interpretability framework linking reservoir dynamics to virtual sources and sensor space, revealing the temporal, spectral, and spatial signatures underlying decoding performance at both the individual and group levels. Together, HeteroRC offers an interpretable approach to decode information from dynamic neural responses, broadening the analytical scope of neural decoding while remaining computationally efficient and free from manual feature engineering, making it particularly suitable for small-sample electrophysiological studies.

## Introduction

Neural decoding has become a central tool in cognitive neuroscience and brain–computer interface research, enabling inference about representational content from patterns of neural activity [1–8]. With the increasing availability of electrophysiological recordings, ranging from non-invasive magnetoencephalography (MEG) and electroencephalography (EEG) to intracranial measures such as local field potentials (LFPs) and multi-unit activity, decoding methods have increasingly exploited the high temporal resolution of these techniques to track neural representations over time. Time-resolved decoding and cross-temporal generalisation are now standard tools for characterising the temporal dynamics of cognitive processes such as perception, attention, memory, and action [6, 9–11]

Despite their broad adoption, most time-resolved decoding studies rely on linear classifiers, typically linear discriminant analysis (LDA) or linear support vector machines (SVM) [6, 9, 12–15]. Comparative benchmarks have shown that, for decoding stimulus-locked and phase-locked information expressed in evoked responses, linear decoders often perform as well as or better than more complex non-linear models [16, 17]. Their robustness, computational efficiency, and interpretability have therefore made linear decoding pipelines, operating on instantaneous signal amplitudes (e.g., voltage for EEG/LFPs or magnetic fields for MEG), the default choice in neural time-series decoding research [6, 18].

However, this methodological standard implicitly favours a restricted class of neural signals: those that are phase-locked to experimental events and expressed as evoked responses [6, 15, 19]. A large body of work indicates that many cognitive variables are instead encoded in neural dynamics that are not phase-locked to stimulus onset, including induced oscillatory power [20–23], phase synchrony [24–27], and scale-free aperiodic activity [28–30]. These dynamics often vary in latency and duration across trials and are therefore poorly captured by conventional decoding approaches that operate independently at each time point on raw instantaneous amplitudes using linear classifiers, which can in turn complicate the interpretation of null decoding findings. In the working memory literature, for example, several studies have reported that memory content cannot be decoded from instantaneous signal traces during delay periods [31–33]. These results have contributed to the proposal that representations may be maintained in an “activity-silent” state [34], while remaining reactivatable by brief external perturbations (i.e., “pinging”) [31, 35, 36]. Importantly, subsequent work has shown that, in some cases, information that is not decodable from raw amplitudes can be recovered from alternative signal features such as induced alpha power [37]. These findings highlight that decoding outcomes depend on the interaction between the neural signal format and the decoding model, and that failures of linear, instantaneous amplitude-based decoding may not uniquely determine absence of task-relevant neural activity, but reflect a mismatch between representational dynamics and decoding assumptions.

While non-phase-locked information can sometimes be decoded by explicitly transforming neural signals into alternative features (e.g., time-frequency representations), such approaches rely on a priori assumptions regarding which signal dimensions carry task-relevant information. In many cognitive paradigms, however, the optimal representational format is unknown and may dynamically evolve across tasks, brain regions, and processing stages. Recent efforts have sought to maximise information recovery by systematically comparing and combining diverse feature sets [38, 39]. Yet, such exhaustive feature engineering remains computationally intensive and demonstrates that integrating multiscale features does not consistently yield additive gains [39]. Alternatively, modern deep learning architectures, such as recurrent neural networks (RNNs) and Transformers, offer a data-driven approach to bypass manual feature extraction, utilising their high non-linear expressivity to learn complex representations directly from raw time series [40]. However, the immense parameter spaces of these fully trainable models inherently demand massive datasets for stable convergence. Consequently, they are susceptible to overfitting and temporal smearing in the small-sample, trial-limited regimes that characterise most cognitive electrophysiology experiments [41].

Addressing these limitations motivates the development of decoding frameworks that operate directly on small-sample neural time series while remaining sensitive to non-linear, non-phase-locked, and multiscale temporal dynamics. Reservoir computing (RC) provides a biologically inspired and computationally efficient approach for mapping time-varying inputs into a high-dimensional dynamic state space using fixed recurrent connectivity and a trained linear readout [42–46]. Through recurrent dynamics, RC models can integrate information over time while preserving interpretability at the readout level, making them attractive for neural decoding applications. However, standard RC implementations implicitly operate at a single intrinsic temporal scale, typically assuming homogeneous time constants across reservoir units. This assumption contrasts with extensive empirical and theoretical evidence that neural activity in cortex spans a hierarchy of intrinsic timescales [47–49], and limits the ability of conventional RC models to represent neural dynamics that unfold concurrently over fast and slow temporal scales.

Here we introduce HeteroRC (Heterogeneous Reservoir Computing), a decoding framework that explicitly incorporates heterogeneous intrinsic time constants into reservoir dynamics. Motivated by empirical and theoretical work demonstrating a hierarchy of intrinsic timescales in cortical activity [47–50], HeteroRC samples reservoir units with a distribution of time constants, enabling the representation of neural dynamics across fast and slow temporal scales within a single recurrent state space. Multichannel neural time series are projected into this high-dimensional dynamic space, and task-relevant information is extracted using a linear readout (e.g., ridge classification). Crucially, the linear readout enables a principled interpretation module, allowing latent reservoir dynamics to be linked back to sensor-level neurophysiological signals. At the individual level, this framework extracts temporal, spectral, and spatial dynamics tailored to single subjects. Furthermore, by utilising a sensor-space matching approach, these findings can be generalised to uncover shared neurophysiological motifs across participants.

Through controlled simulations, we first demonstrate that HeteroRC significantly outperforms conventional linear decoders and a suite of artificial neural networks (ANNs; including RNNs, LSTMs, Transformers, and EEGNet) on evoked responses, while robustly capturing induced oscillatory power, phase synchrony, and aperiodic modulations—neural dynamics that are typically inaccessible to conventional linear methods. Applying the method to an empirical motor imagery neuroimaging dataset [51], we demonstrate improved decoding performance relative to conventional linear decoders, particularly during internally generated imagery periods. Moreover, HeteroRC supports robust cross-temporal generalisation, capturing systematic changes in representational dynamics from cue-driven to internally-maintained states. Leveraging the interpretation framework, we identify the temporal, spectral, and spatial characteristics of neural signal components that contribute most strongly to successful decoding at the individual-subject level, and further reconstruct grand-average virtual sources to confirm that these distinct mechanistic signatures are robustly conserved across the group. Furthermore, using an attentional priority mapping dataset [36], we show that latent information not readily decodable from instantaneous amplitude-based decoding can be recovered using HeteroRC, and demonstrate that decoding in perturbed and unperturbed conditions relies on partially distinct neural signal components, revealing different encoding regimes despite shared representational content. Together, these results establish HeteroRC as a robust and interpretable framework for neural decoding, enabling latent information in dynamic brain signals to be recovered and systematically interpreted beyond conventional approaches.

## Results

### HeteroRC

To overcome the limitations of conventional linear decoders in capturing non-phase-locked and nonlinear neural dynamics, while also avoiding the requirement for large training datasets characteristic of deep learning architectures, we developed HeteroRC. HeteroRC is a decoding framework that projects multichannel neural time series into a high-dimensional recurrent state space governed by heterogeneous intrinsic time constants (Figure 1a). These time constants are sampled from a log-normal distribution, establishing a multiscale temporal filter bank that enables simultaneous integration of fast, transient responses and slower, persistent neural dynamics. This allows HeteroRC to operate directly on raw neural time series without requiring explicit feature engineering (e.g., pre-computed spectral power or phase), while retaining the computational efficiency of a fixed reservoir coupled with a trained linear readout.

**Figure 1.**
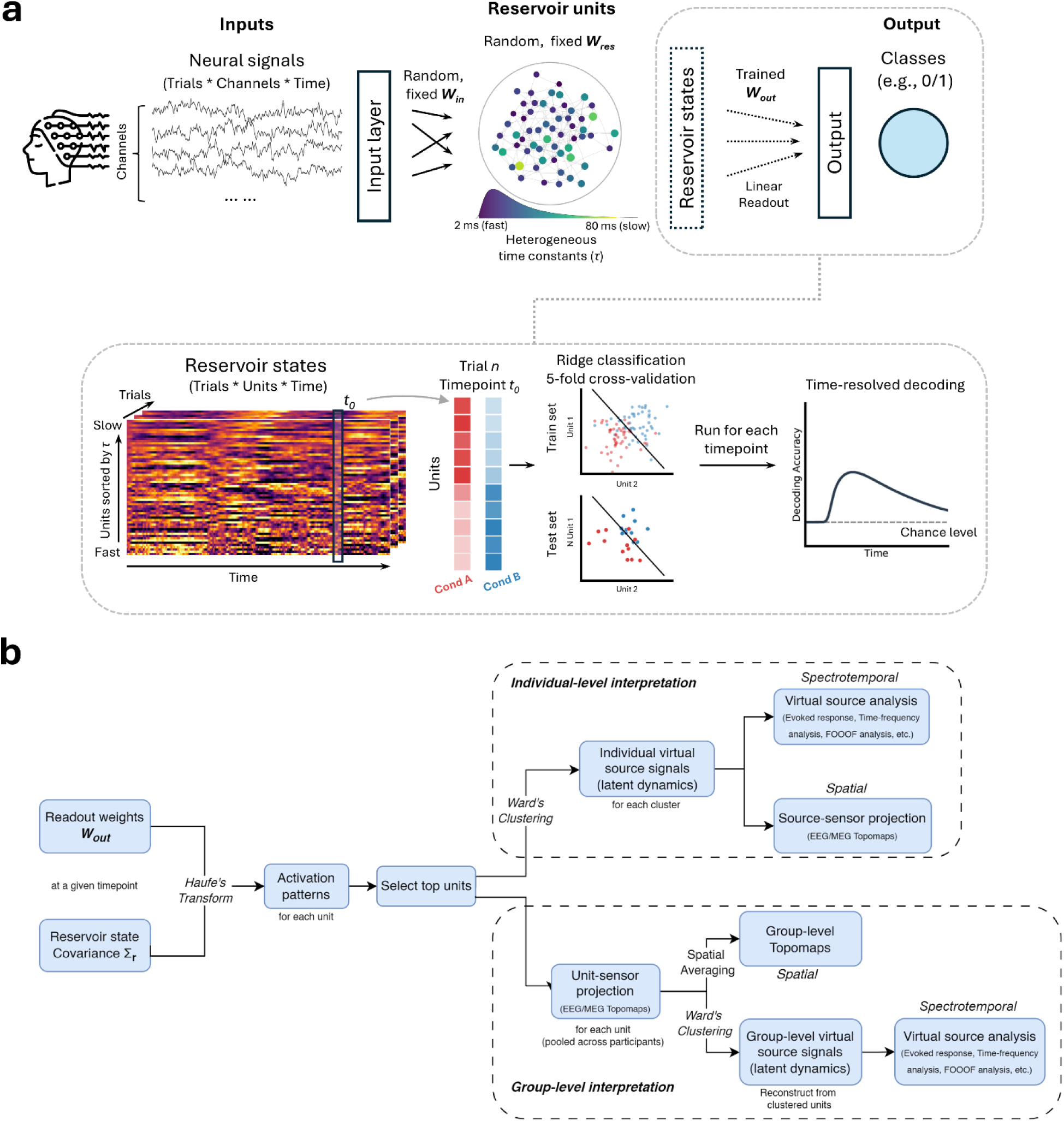
HeteroRC decoding and interpretation framework. (a) Decoding framework. Multichannel neural signals (trials × channels × time) are provided as inputs to a recurrent reservoir. Inputs are linearly projected to reservoir units through a fixed, randomly initialised input weight matrix (*W*_in_), with full connectivity between input channels and reservoir units. Reservoir units are connected via a fixed recurrent weight matrix (*W*_res_) with sparse random connectivity (10% non-zero connections), scaled to ensure stable echo-state dynamics. Each reservoir unit is endowed with an intrinsic time constant (*τ*), sampled from a log-normal distribution, resulting in heterogeneous temporal integration properties spanning fast and slow timescales. Neural inputs are thus transformed into high-dimensional reservoir state trajectories that capture multiscale temporal dynamics. At each time point, reservoir states are decoded using a trained linear readout (*W*_out_ ; ridge classification), yielding time-resolved decoding performance for task conditions or stimulus classes. (b) Interpretation framework. Trained readout weights are first projected back into reservoir space using a covariance-based activation pattern analysis (Haufe’s transform), identifying reservoir units that contribute most strongly to decoding. From here, the framework branches into two analytical tracks. For individual-level interpretation, the dynamics of informative units are directly clustered to extract individual-specific latent virtual source signals. These virtual sources are analysed in the time and frequency domains (e.g., evoked responses, time-frequency representations, fitting oscillations & one over (FOOOF) analysis) and further projected back to sensor space to reveal their corresponding spatial topographies, enabling joint temporal, spectral, and spatial interpretation of decoding performance. For group-level interpretation, selected units from all participants are first projected to a common sensor space. These spatial topographies are pooled and globally clustered, and the resulting cluster assignments are mapped back to individual reservoirs to reconstruct group-level virtual source signals. These sources are subsequently subjected to the same temporal, spectral, and spatial analyses mentioned above.

To address the interpretability challenges associated with recurrent models, we developed an interpretation framework that links reservoir dynamics to physiological signal generators (Figure 1b). First, we apply a covariance-based activation pattern analysis [18] to project the learned readout weights back into reservoir space and identify units that contribute most strongly to decoding. By clustering the temporal dynamics of these informative units, we extract latent virtual source signals that capture task-relevant dynamics within the reservoir. These virtual sources can be analysed directly in the time and frequency domains and are further projected back to sensor space to reveal their corresponding spatial topographies. Together, this framework disentangles the temporal, spectral, and spatial signatures underlying decoding performance, allowing phase-locked evoked responses to be distinguished from induced oscillatory and aperiodic components, and directly linking decoding outcomes to interpretable neural signal mechanisms.

### HeteroRC decodes diverse classes of neural dynamics under controlled simulations

To systematically evaluate the sensitivity of HeteroRC to distinct classes of neural dynamics, we generated synthetic datasets simulating five canonical signal types commonly observed in neural time series recordings: 1) phase-locked evoked responses, 2) induced oscillatory power, 3) inter-site phase clustering (ISPC), and 4) slope and 5) offset of aperiodic spectral modulations. Simulated signals were embedded in realistic 1/*f* background activity with white noise over 0.8-s epochs, with task-relevant modulations confined to a 0.2–0.6-s time window and subject to trial-by-trial temporal jitter (see Methods). For simulations involving oscillatory power and phase synchrony, task-relevant modulations were centred at 10 Hz, reflecting the canonical alpha-band activity; however, identical decoding patterns were obtained when modulations were placed at other frequencies (Figure S1). Crucially, the strength of each signal type was manipulated independently while holding the remaining components constant, ensuring that decoding performance reflected sensitivity to the targeted dynamical feature. We employed an event-related design with 5-fold cross-validation, where feature scaling and model training were strictly separated within each fold to prevent data leakage.

We first evaluated decoding performance in simulations where discriminative information was carried by phase-locked evoked responses, a regime typically considered the optimal use case for linear, instantaneous amplitude-based classifiers. In this setting, HeteroRC outperformed both LDA and SVM, achieving higher decoding accuracy within the task-relevant window while preserving accurate temporal localisation of the evoked response peak (Figure 2a).

**Figure 2.**
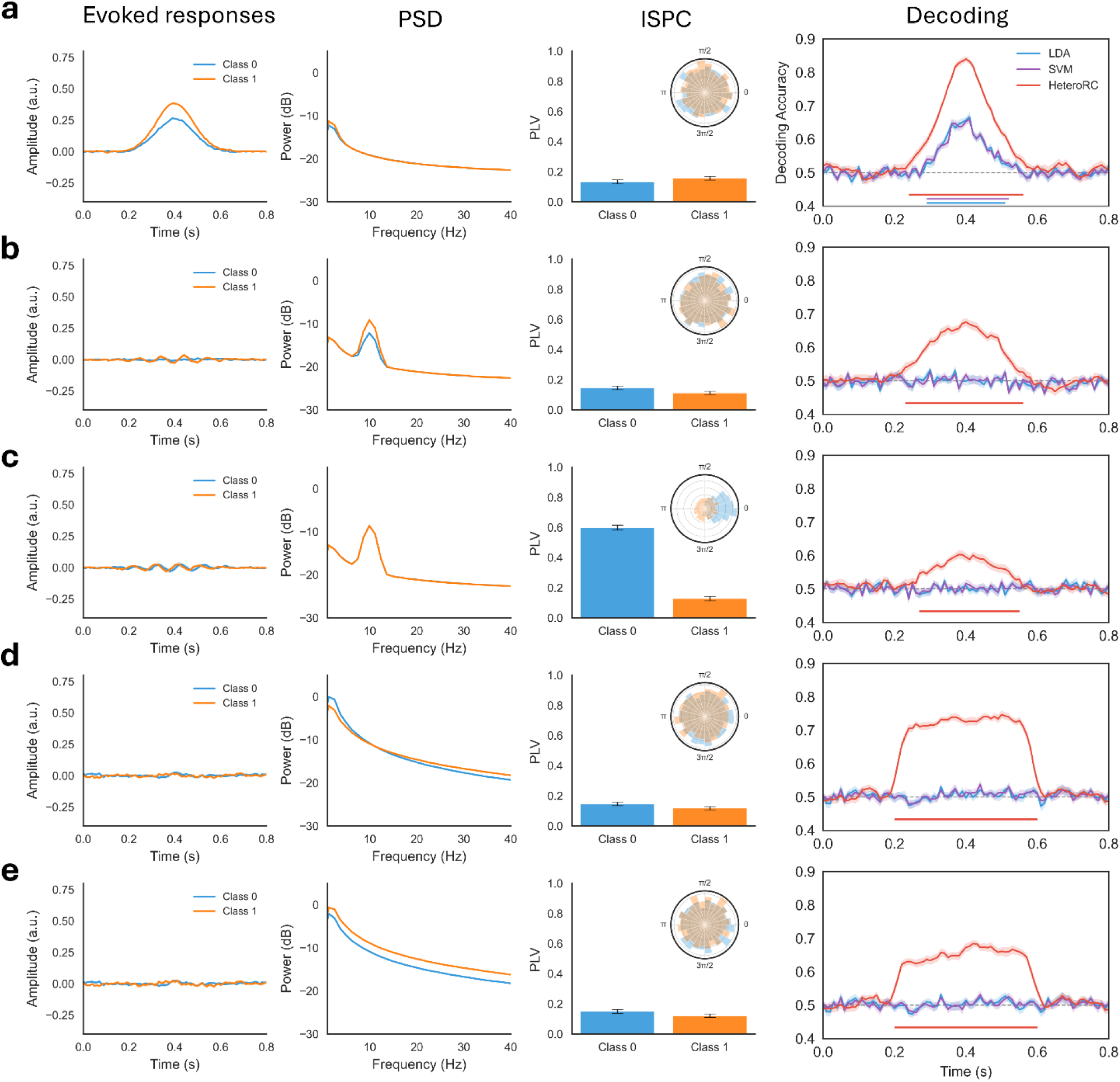
HeteroRC decodes diverse classes of neural dynamics in controlled simulations. Simulated datasets were generated to isolate and test decoding sensitivity to five distinct classes of neural dynamics: phase-locked evoked responses (a), induced oscillatory power with randomised phase (b), inter-site phase clustering (ISPC) (c), and aperiodic spectral modulations affecting the 1/*f* slope (d) and offset (e). In all conditions, task-relevant modulations were confined to a ∼0.2–0.6-s time window and subject to trial-by-trial temporal jitter. For simulations involving oscillatory power and ISPC, task-relevant modulations were centred at 10 Hz (for results at different frequencies see Figure S1). For each simulation, the left panels illustrate the underlying signal differences between conditions, including evoked responses (time domain), power spectral density (PSD, frequency domain), and ISPC (phase synchrony), while polar plots depict phase distributions. Right panels show time-resolved decoding accuracy for HeteroRC (red) compared with conventional linear decoders (LDA, blue; SVM, purple). Horizontal bars indicate time points with decoding performance significantly above chance (*p* < 0.05, corrected by cluster-based permutation test).

A pronounced divergence in performance between HeteroRC, LDA and SVM emerged when discriminative information was embedded in non-phase-locked dynamics, including induced oscillatory power with randomised phase, ISPC, and aperiodic slope and intercept modulations. Under these conditions, both LDA and SVM applied to raw time series failed to recover task-relevant information, yielding performance near chance levels (Figure 2b–d). In contrast, HeteroRC robustly decoded these latent dynamics, despite the absence of consistent phase-locked amplitude differences. It moreover accurately recovered the time-window in which the latent dynamics had been applied.

To explicitly contrast HeteroRC with conventional feature engineering pipelines, we additionally evaluated the performance of LDA when it was trained and tested on data that was manually transformed into time-frequency space – specifically, on 8–12 Hz alpha-band power extracted via Morlet wavelets and a Hilbert filter approach (Figure S2). We again tested this approach for detecting underlying effects embedded in evoked responses, induced oscillatory power with randomised phase, ISPC, and aperiodic slope and intercept modulations. As expected, the feature engineering approach successfully decoded information when the underlying neural dynamics matched the features of the data that the manual engineering approach targeted. Thus, LDA based on 8-12Hz alpha-band power was able to discriminate the classes when the underlying signal varied in 10 Hz induced power, and when the underlying modulation was a change in the broadband aperiodic intercept, since this manipulation also altered absolute alpha-band power. It could also weakly and partially detect the underlying modulation of aperiodic slope, for the same reason. However, the engineering pipeline completely failed to recover phase-locked evoked responses, ISPC, and induced power outside the pre-specified filter range (e.g., 20 Hz). Furthermore, even when decoding was successful (e.g., for 10 Hz power), both time-frequency methods, and particularly the wavelet approach, exhibited pronounced temporal smearing extending beyond the ground-truth 0.2–0.6-s window. These results demonstrate that explicit feature extraction is not only bottlenecked by manual parameterisation but also susceptible to integration-induced temporal blurring.

Given this observation, and because HeteroRC also inherently relies on its own recurrent temporal integration, an important concern is whether its robust decoding of latent dynamics could be similarly confounded by temporal smearing or systematic latency shifts. To address this, we compared different reservoir processing strategies in controlled simulations, including unidirectional processing and alternative bidirectional fusion schemes (see Methods for details). As shown in Figure S3, in simulations dominated by evoked responses and aperiodic activity, unidirectional reservoirs exhibited systematic temporal lags and broadened decoding peaks, consistent with integration-induced smearing. Bidirectional processing substantially reduced these effects by cancelling direction-specific temporal biases. Among bidirectional strategies, multiplicative fusion (the method used in Figure 2) provided sharper temporal localisation than averaging (as in Figure S3), while preserving robust decoding performance. These results indicate that, unlike conventional filtering approaches, the sustained decoding observed with HeteroRC more faithfully reflects true underlying neural dynamics rather than algorithmic lag, thereby motivating the use of bidirectional multiplicative fusion throughout the main analyses.

Together, these simulations demonstrate that HeteroRC not only exceeds the performance of conventional linear decoders in their preferred regime of phase-locked evoked responses, but it also generalizes to reliably recover information encoded in non-phase-locked and aperiodic neural dynamics. This broad sensitivity enables accurate, time-resolved decoding across a wide range of biologically meaningful signal formats commonly present in neural time series recordings.

### HeteroRC outperforms artificial neural networks in decoding evoked responses

While ANNs offer high nonlinear expressivity, their effectiveness inherently scales with the availability of large training datasets. To contextualise HeteroRC against this class of machine learning approaches, we compared its performance against four canonical ANNs: RNNs [52] and Long Short-Term Memory (LSTM) networks [53], which rely on recurrent hidden states to integrate temporal sequences; Transformers [54], which use self-attention mechanisms capture global temporal dependencies; and EEGNet [55], a convolutional neural network (CNN)-based architecture specifically optimised for the spatial and temporal constraints of EEG signals. We focused this comparative benchmark specifically on phase-locked evoked responses, as these highly consistent signals represent the most straightforward scenario for ANNs to achieve stable performance. Due to its convolutional architecture, EEGNet was evaluated exclusively using a windowed formulation to generate a time-resolved decoding profile comparable to HeteroRC. In contrast, the other models were assessed in both standard sequence-to-sequence and windowed regimes.

For the general-purpose ANNs (RNN, LSTM, and Transformer), we first evaluated them in a standard sequence-to-sequence regime. In this approach, the models process the full temporal epoch at once and are trained to output a continuous trajectory of class predictions, generating a discrete decision at every individual time step. For the RNN and LSTM, this prediction relies on the accumulated signal history, whereas the Transformer globally integrates both past and future context across the entire epoch. In this setting, HeteroRC substantially outperformed these models (Figure 3a). Notably, RNN and LSTM exhibited a pronounced temporal lag and smearing, with their decoding accuracy peaking later but persisting longer than the ground-truth neural modulation. This reflects the integration time required to accumulate evidence from past inputs and the persistence of information within their hidden states even after the underlying signal had ceased. The standard Transformer poorly localized neural dynamics in time, as its unmasked global self-attention mechanism integrates both past and future context across the entire epoch.

**Figure 3.**
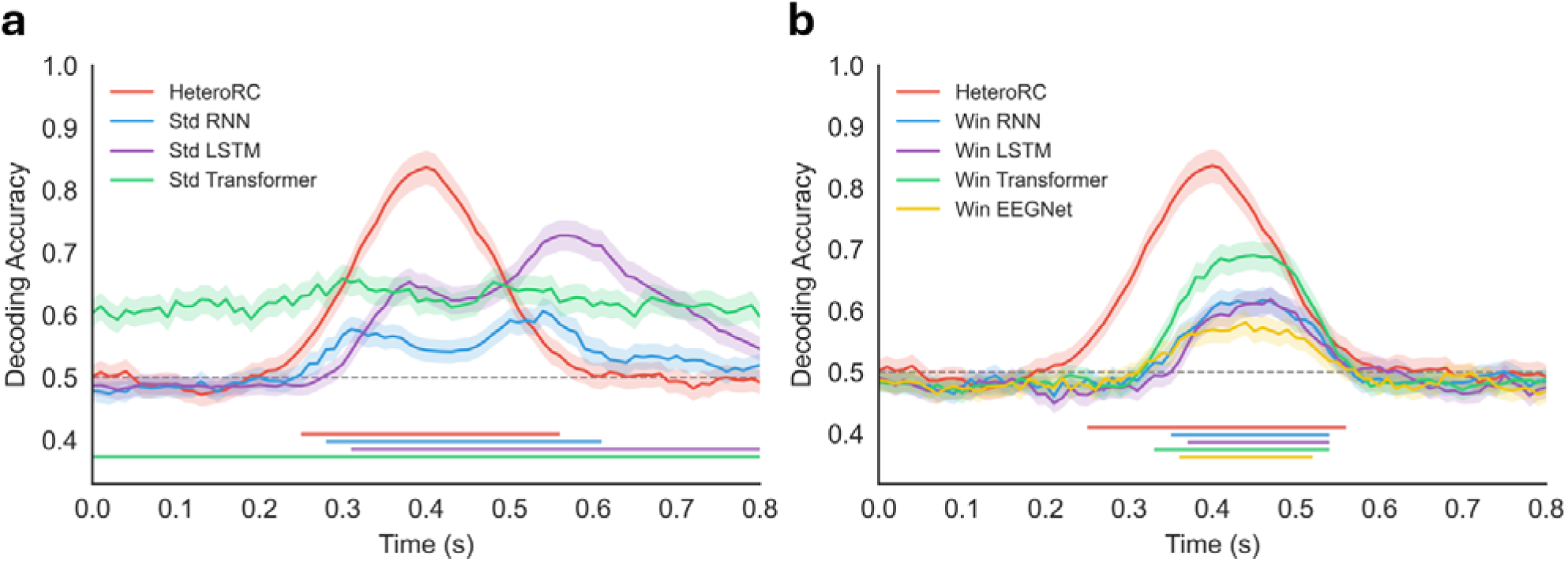
HeteroRC outperforms ANNs in decoding evoked responses. Time-resolved decoding accuracy for simulated phase-locked evoked responses, comparing HeteroRC (red) against standard and windowed ANNs. Task-relevant modulations were confined to a ∼0.2–0.6-s time window. (a) Comparison with standard sequence-to-sequence models: Recurrent Neural Network (Std RNN, blue), Long Short-Term Memory network (Std LSTM, purple), and Transformer (Std Transformer, green). (b) Comparison with windowed (sequence-to-one) models utilising a 100-ms sliding window, including Windowed RNN (blue), LSTM (purple), Transformer (green), and the EEGNet (yellow). EEGNet was evaluated exclusively using the windowed formulation to respect its convolutional architecture. Shaded regions denote the standard error of the mean across simulated subjects. Horizontal dashed lines indicate chance-level performance (0.5). Solid horizontal bars at the bottom indicate time windows with robust decoding performance for the corresponding models.

To mitigate these issues and provide a more rigorous baseline, we next trained windowed variants of the ANNs alongside EEGNet. By applying a 100ms sliding window across the epoch, we strictly bounded the temporal integration horizon, forcing the models to rely on local signal features rather than accumulated history or future context. Although this windowing strategy improved the temporal localization of the decoding results, partially correcting the temporal lag and smearing, HeteroRC continued to demonstrate superior decoding accuracy and temporal precision (Figure 3b).

These results indicate that HeteroRC, utilising a fixed reservoir with heterogeneous intrinsic time constants, captures multiscale neural dynamics more efficiently than fully trainable ANNs when data is limited. By integrating these dynamics with bidirectional multiplicative fusion to actively cancel temporal lags, HeteroRC achieves a highly favourable trade-off between representational richness, data efficiency, and temporal precision.

### HeteroRC decodes sustained internally generated motor imagery representations

To validate HeteroRC on empirical data, we first applied the framework to a motor imagery dataset (*N* = 9; BCI Competition IV 2a [51]), which requires decoding of four classes of imagined movements (left hand, right hand, feet, and tongue) from a 22-channel EEG (Figure 4a). Motor imagery relies on internally generated neural activity with variable onset and duration across trials, providing a stringent test for decoding non-phase-locked dynamics. In addition, training and evaluation data were collected in separate recording sessions on different days for each participant, requiring models to generalise across sessions despite potential non-stationarities in signal quality or electrode impedance. Given that our controlled simulations demonstrated near-identical performance between LDA and linear SVM, we restricted our subsequent comparisons to LDA for computational efficiency.

**Figure 4.**
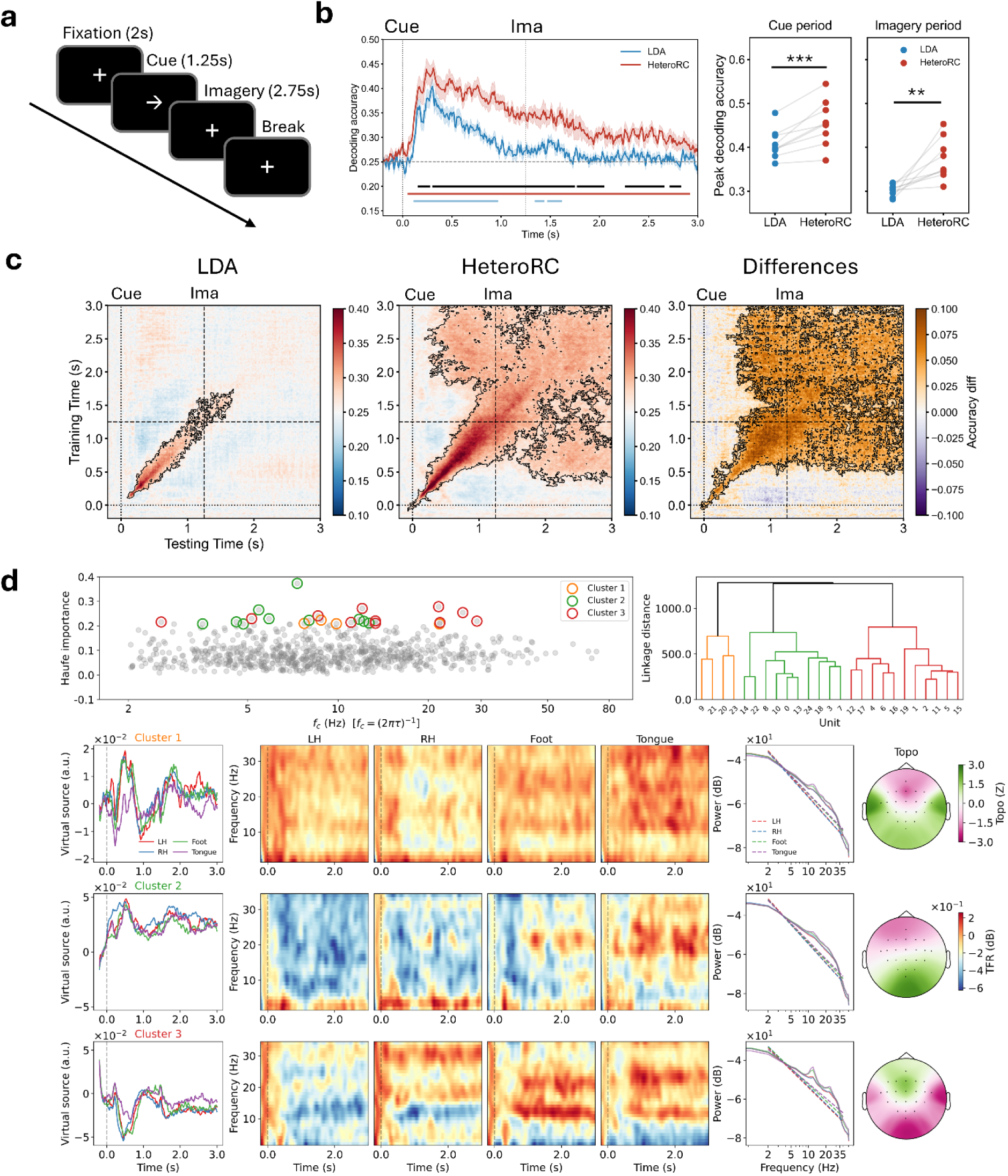
HeteroRC decodes internally generated motor imagery and captures representational dynamics. (a) Experimental design of the motor imagery task [51]. Each trial consisted of a fixation period, followed by a visual cue (an arrow pointing left, right, down, or up) indicating the movement to be imagined (left hand (LH), right hand (RH), feet, or tongue, respectively), and a sustained motor imagery period in the absence of external sensory input. (b) Left: Time-resolved decoding accuracy over time for HeteroRC (red) and linear discriminant analysis (LDA; blue). Shaded regions denote the standard error of the mean across participants. Horizontal bars indicate time points with decoding accuracy significantly above chance (*p* < 0.05, corrected by cluster-based permutation test) for HeteroRC (red), LDA (blue) and the difference between them (black). Right: Peak decoding accuracy for each participant during the cue period and the sustained imagery period. Each dot represents one participant, with lines connecting LDA and HeteroRC within subjects. HeteroRC shows significantly higher peak decoding accuracy than LDA in both periods (\*\**p* < 0.01; \*\*\**p* < 0.001). (c) Cross-temporal generalisation matrices. Decoding accuracy as a function of training time (y-axis) and testing time (x-axis) for LDA (left), HeteroRC (middle), and their difference (HeteroRC minus LDA, right). Black contours indicate regions significantly above chance (*p* < 0.05, corrected by cluster-based permutation test). (d) Interpretation of reservoir dynamics during sustained imagery for an individual participant. Top: Haufe-transformed activation patterns plotted as a function of reservoir unit intrinsic timescale, with most 25 informative units grouped into three clusters using hierarchical clustering. Bottom: Analyses of virtual source signals derived for each cluster, showing trial-averaged temporal profiles, time–frequency representations, power spectral density estimates of oscillatory and aperiodic components (derived using FOOOF), and corresponding sensor-space projections.

We first evaluated time-resolved decoding accuracy, comparing HeteroRC against LDA (Figure 4b). In the experimental paradigm, each trial consisted of a fixation period, followed by a visual cue indicating the movement to be imagined, and a sustained motor imagery period in the absence of external sensory input. Following cue onset, both HeteroRC and LDA showed a rapid increase in decoding accuracy with comparable latencies. However, LDA performance declined to near-baseline levels before cue offset, whereas HeteroRC maintained robust and significant decoding accuracy. Crucially, during the sustained imagery period (from 1.25 s onward), when no external stimulus was present and decoding depended entirely on internally maintained representations, HeteroRC significantly outperformed the linear decoder. HeteroRC sustained above-chance decoding for more than 1.5 s, whereas LDA exhibited reliable decoding for only approximately 250 ms. In addition, inspection of individual-subject decoding dynamics (Figure S4) suggested that HeteroRC numerically outperformed LDA in decoding accuracy for all participants. To quantify these individual-subject level effects, we compared peak decoding accuracy between HeteroRC and LDA, separately for the cue period (0–1.25 s) and the sustained imagery period (1.25–3 s). HeteroRC showed significantly higher peak decoding accuracy than LDA in both cue (*t*(8)= 5.32, *p* < 0.001) and imagery periods (*t*(8) = 3.91, *p* = 0.004) (Figure 4b, right). This indicates that the sustained decoding advantage of HeteroRC is robust across participants and not driven by a small subset of subjects.

To further characterise the temporal structure of the underlying neural codes, we computed cross-temporal generalisation matrices (Figure. 3c). The linear decoder exhibited a predominantly diagonal generalisation pattern confined to the cue period and the early imagery phase, indicating reliance on transient, time-specific and likely phase-locked features. In contrast, HeteroRC revealed a dynamic evolution in representational structure. During the early cue phase (before ∼600 ms), generalisation was largely diagonal, consistent with dynamically evolving sensory representations. This pattern progressively transitioned into a broad, square-like generalisation structure during the late cue phase (after ∼600 ms) and persisted throughout the sustained imagery period, indicating the emergence of a temporally stable representational regime.

To identify the neural mechanisms supporting this sustained decoding during imagery, we applied the interpretation framework to an individual participant (Subject 1) at the peak decoding time point within the imagery period (2.3 s). Using a covariance-based activation pattern analysis [18], we identified the top 25 most informative reservoir units and grouped them into three functional clusters via hierarchical clustering, yielding three virtual source signals that captured distinct latent dynamics contributing to decoding (see Methods for details). Inspection of these virtual sources revealed that HeteroRC leveraged multiple signal characteristics to distinguish imagined motor commands. Some components (e.g., Cluster 1) exhibited distinct condition-specific temporal modulations, whereas some reflected broader spectral dynamics (Cluster 2). Notably, Cluster 3 captured differences in oscillatory power within the alpha/mu range (8-13 Hz) as well as class-dependent variations in aperiodic spectral slope, which differentiated hand imagery from foot and tongue imagery. Projection of these sources back to sensor space further revealed spatially distinct cortical topographies associated with each dynamical component. Together, these results indicate that the reservoir integrates information across temporal, spectral, and spatial domains to form a robust representation of internally generated motor states.

In summary, HeteroRC robustly decodes internally generated motor imagery representations that are difficult to recover using conventional linear models. Beyond improved decoding accuracy, cross-temporal generalisation analyses demonstrate that HeteroRC tracks the transformation of neural representations from transient cue-driven activity to stable, internally maintained imagery states. The accompanying interpretation framework further reveals the distinct temporal, spectral, and spatial signal components supporting this performance, establishing HeteroRC as a powerful and interpretable tool for characterising cognitive processes that unfold in the absence of external sensory input.

### HeteroRC uncovered latent attentional priority representations invisible to linear decoding

Finally, we evaluated whether HeteroRC could recover task-relevant information from neural states that are typically considered inaccessible to standard raw amplitude-based decoding approaches. We addressed this question using an attentional priority mapping dataset (*N* = 23, Figure 5a) [36] in which participants acquired spatial priority maps through statistical learning. Briefly, participants learnt implicitly that one of eight locations in a visual display was more likely to contain a target (a singleton shape). Previous analyses using linear classifiers reported that neural representations of these priority locations was undetectable during the inter-trial interval and could only be detecting following a brief visual impulse (“ping”), leading to the proposal that the representations were maintained in an activity-silent state.

**Figure 5.**
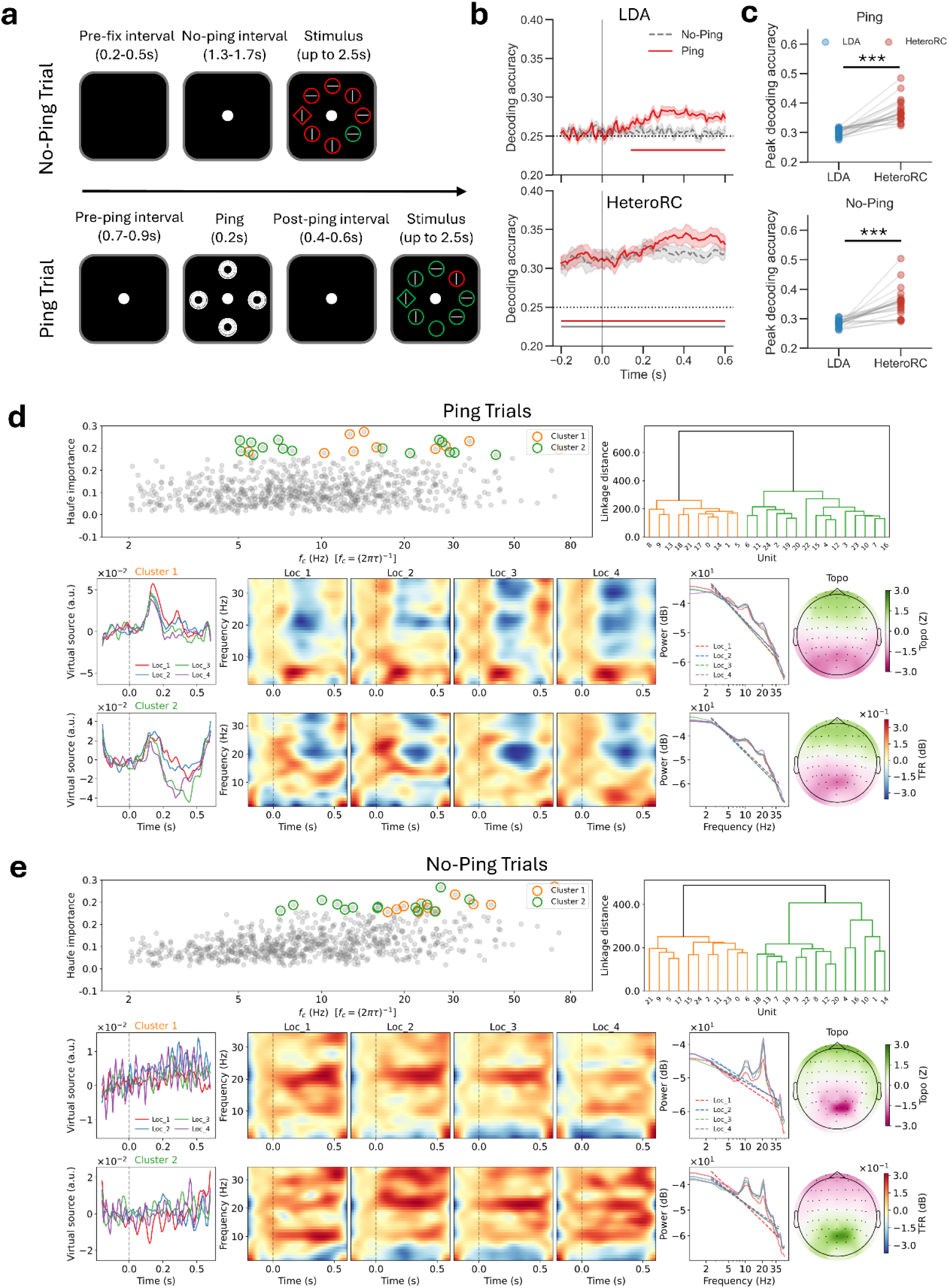
HeteroRC recovers latent task information in an attentional priority mapping task. (a) Experimental design of the attentional priority mapping task [36]. Participants had to report the orientation of a line surrounded by a singleton shape. Participants implicitly learned spatial regularities in target locations across blocks, forming a learned attentional priority map. In Ping trials, a brief visual impulse (200 ms) was presented during the inter-trial interval. In No-Ping trials, no visual stimulus was presented, but a matched trigger event was recorded at an equivalent time point. The subsequent visual search display required participants to report the target feature and was not used for decoding. (b) Time-resolved decoding accuracy of spatial priority for linear discriminant analysis (LDA; top) and HeteroRC (bottom) in Ping (red) and No-Ping (grey dashed) conditions, aligned to ping (or no-ping interval) onset. Shaded regions denote the standard error of the mean across participants. Horizontal bars indicate time points with decoding accuracy significantly above chance (*p* < 0.05, corrected by cluster-based permutation test). The LDA results replicate the results reported in the original paper, in which spatial priority can only be decoded in the ping (red) condition. However, HeteroRC shows sustained decoding of attentional priority before and throughout this time window, for both ping (red) and no-ping (grey). (c) Peak decoding accuracy for each participant in Ping (top) and No-Ping (bottom) conditions. Each dot represents one participant, with lines connecting LDA and HeteroRC within subjects. HeteroRC shows significantly higher peak decoding accuracy than LDA in both conditions (\*\*\**p* < 0.001). (d) Interpretation of reservoir dynamics in Ping trials for an individual participant. Top: Haufe-transformed activation patterns plotted as a function of reservoir unit intrinsic timescale, with most 25 informative units grouped into two clusters using hierarchical clustering. Bottom: Analyses of virtual source signals derived for each cluster, showing trial-averaged temporal profiles, time–frequency representations, power spectral density estimates of oscillatory and aperiodic components (derived using FOOOF), and corresponding sensor-space projections. These clusters primarily capture distinct phase-locked evoked responses. (e) Interpretation of reservoir dynamics in No-Ping trials (same analyses as in d). In contrast, these clusters capture non-phase-locked induced alpha/beta dynamics and aperiodic shifts.

We compared the decoding of spatial priority between HeteroRC and LDA in both Ping and No-Ping conditions (Figure 5b). Consistent with the original publication [36], LDA successfully decoded spatial priority following the visual impulse but failed to achieve above-chance performance in the absence of the impulse, suggesting the absence of decodable information during No-Ping trials. In contrast, using HeteroRC, spatial priority information was robustly decoded in both Ping and No-Ping conditions. There was significant decoding accuracy observed throughout the epoch in both cases, including before the ping onset (and equivalent no-ping timepoint). Nonetheless, the ping still elicited a transient numerical increase in decoding accuracy.

At the individual-subject level, again, HeteroRC tended to show higher decoding accuracy than LDA for nearly all participants in both Ping and No-Ping conditions, as illustrated by per-subject time-resolved decoding results (Figure S5). To quantify these effects across participants, we compared peak decoding accuracy between HeteroRC and LDA separately for Ping (*t*(22) = 8.92, *p* < 0.001) and No-Ping (*t*(22) = 6.28, *p* < 0.001) trials. HeteroRC showed significantly higher peak decoding accuracy than LDA in Ping and No-Ping conditions (Figure 5c).

To identify the neural features supporting this latent information, we applied the interpretation framework to an individual participant (Subject 24) at the peak decoding time point for Ping (0.34 s) and No-Ping (0.48 s) trials. In Ping trials (Figure 5d), virtual source analysis revealed prominent evoked responses that discriminated priority locations (both clusters), consistent with phase-locked reactivation of the priority map by the external impulse. This involved spatially distributed generators across frontal and posterior regions. In contrast, No-Ping trials (Figure 5e, both clusters) showed no discernible evoked components. Instead, decoding was driven by induced neural dynamics: virtual sources exhibited condition-specific oscillatory activity in the alpha (∼10 Hz) and beta (∼20 Hz) bands, accompanied by shifts in aperiodic spectral slope and offset. Projection of these sources to sensor space localised the dominant contributions primarily to posterior cortical regions.

These results suggest that neural information traditionally considered inaccessible to instantaneous amplitude-based linear decoding (e.g., attentional priority information) may in fact be continuously maintained in latent, non-phase-locked dynamics, which can be readily recovered by HeteroRC. On the other hand, the accompanying interpretation framework further reveals that decoding in Ping and No-Ping conditions is supported by distinct neural signal components. This observation adds important new insight to the debate surrounding activity-silent mechanisms, suggesting *both* that previously “hidden” information may in fact have been maintained in non-phase locked activity, and also that additional evoked activity, more closely aligned with activity-silent mechanisms, can be recovered through pinging. These observations showcase the additional insight possible through the proposed interpretability approach using HeteroRC.

### Group-level interpretation of latent dynamics via sensor-space matching

While the single-subject interpretation above provides highly resolved spatiotemporal signatures, a fundamental challenge in applying reservoir computing to neural data is generalising these mechanistic motifs across groups of participants. Because the randomly initialised latent reservoir spaces are idiosyncratic to each participant, direct cross-subject comparison of internal reservoir units is challenging. To overcome this, we developed a sensor-space matching approach. We extracted the top 10 most informative units from each participant and projected their activity back to the standardised EEG sensor level, a physical space that is anatomically comparable across individuals. We then performed a global clustering analysis on the spatial topographies of all extracted units, grouping them into distinct spatial clusters representing shared neurophysiological motifs.

By mapping these spatially matched units back to their respective reservoirs, we reconstructed grand-average virtual source signals across participants (Figure 6). Applying this framework first to the motor imagery dataset, the group-level interpretation isolated representational motifs across the cohort that complemented the highly specific dynamics observed at the individual level. Specifically, the global spatial clustering disentangled three distinct topographical components: two lateralised clusters peaking over the left (Cluster 1) and right (Cluster 3) central electrodes, and a centrally distributed cluster over the midline (Cluster 2). These distinct spatial topographies cleanly indicated lateralised hand imagery from centrally represented foot and tongue imagery. Furthermore, analysis of the reconstructed virtual sources revealed that these spatial clusters captured functionally distinct dynamical profiles. Consistent with the single-subject observations, some components (e.g., Cluster 2) exhibited more pronounced condition-specific temporal modulations, whereas the lateralised components (Clusters 1 and 3) were characterised by strong, oscillatory power modulation in the alpha/mu and beta bands.

**Figure 6.**
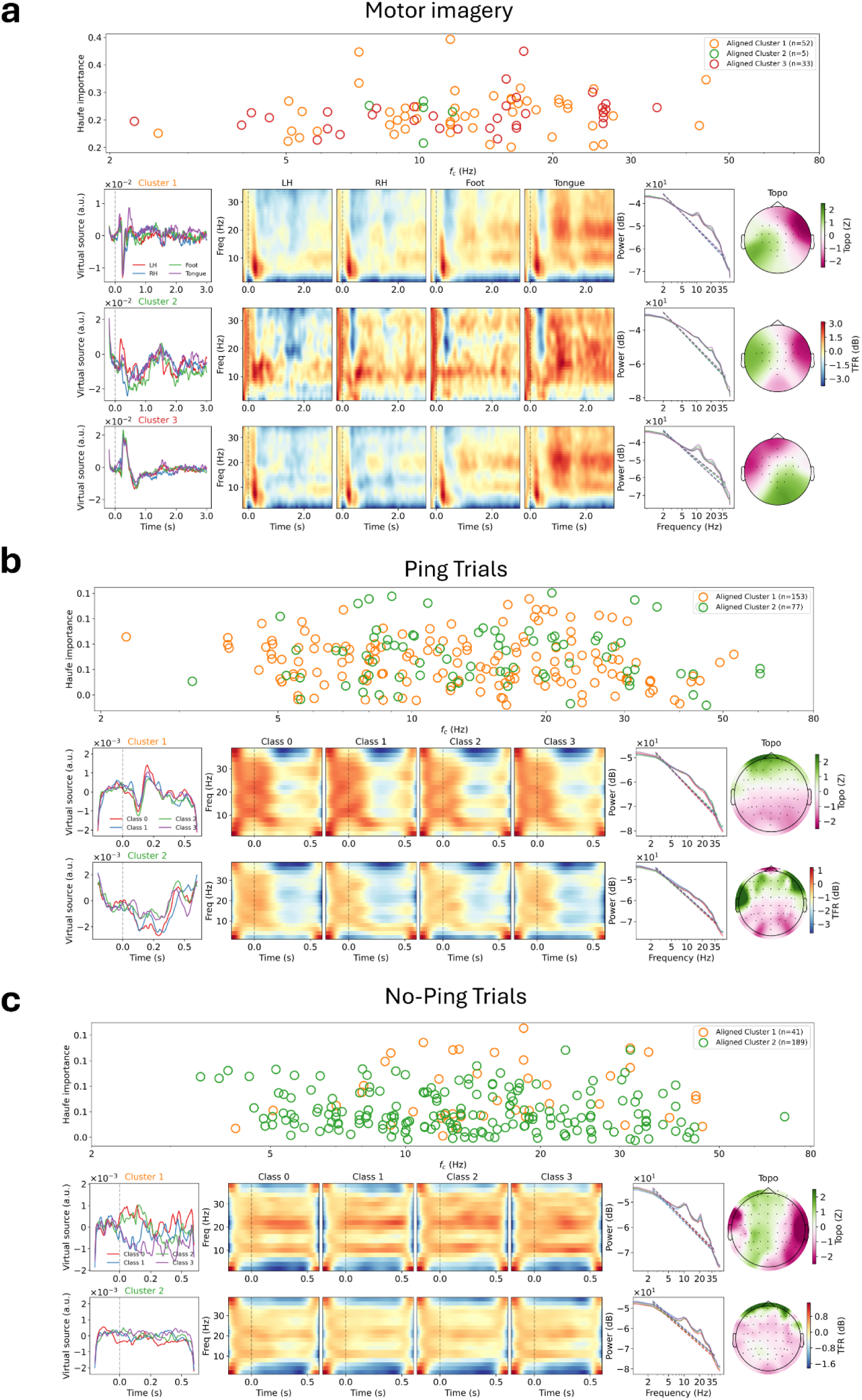
Group-level interpretation of latent dynamics via sensor-space matching. (a) Experimental Group-level interpretation of reservoir dynamics in the motor imagery dataset across all participants. Top: Haufe-transformed activation patterns of the top 10 most informative units per participant plotted as a function of reservoir unit intrinsic timescale, grouped into three global clusters based on their spatial topographies. Bottom: Analyses of grand-average virtual source signals reconstructed for each spatial cluster, showing trial-averaged temporal profiles, time–frequency representations, power spectral density estimates of oscillatory and aperiodic components (derived using FOOOF), and corresponding grand-average sensor-space projections. (b) Group-level interpretation of reservoir dynamics in Ping trials from the attentional priority dataset (same analyses as in a, grouped into two global clusters). (c) Group-level interpretation of reservoir dynamics in No-Ping trials.

Similarly, in the attentional priority dataset, the group-level virtual sources corroborated the mechanistic divergence between maintenance states seen in the example individual subject level above. Decoding in the Ping condition was predominantly driven by strong, phase-locked evoked responses localised to posterior sensors (Figure 6b). In contrast, decoding in the No-Ping condition reflected induced oscillatory power modulations (Figure 6c).

Collectively, these findings not only reinforce that successful decoding of neural representations relies on the joint contribution of diverse temporal, spectral, and spatial neural signatures, but also demonstrate that HeteroRC can reliably extract and disentangle heterogeneous neural codes across individuals without requiring direct hyperalignment of their idiosyncratic latent state spaces.

## Discussion

Neural decoding is widely used to infer when and how information is represented in the brain, particularly in time-resolved analyses of EEG, MEG, and LFPs signals [2, 6, 9]. Conventional decoding pipelines in neuroscience predominantly rely on linear classifiers applied to sensor-level raw amplitude time series, which are effective for capturing stimulus-locked activity [16, 17] but are less sensitive to information encoded in non-phase-locked or nonlinear neural dynamics [21, 37]. As a consequence, a failure to decode information using standard pipelines risks being misinterpreted as an absence of active neural coding. Our findings caution against this, demonstrating that decodability is not solely a property of the neural representation itself, but depends critically on the interaction between neural dynamics and the assumptions of the decoding model. Across controlled simulations and two empirical datasets, HeteroRC consistently recovered task-relevant information that was inaccessible to conventional linear amplitude-based decoders, including information encoded in induced oscillatory activity, phase synchronisation, aperiodic dynamics, and internally generated neural states. Crucially, the accompanying individual- and group-level interpretation framework enabled these decoding results to be linked back to identifiable temporal, spectral, and spatial neural signatures, revealing distinct dynamical regimes supporting decoding under different task conditions.

A key feature of the present decoding framework is its ability to operate directly on raw neural time-series data without explicit feature engineering, while remaining sensitive to information expressed in both phase-locked and non-phase-locked and nonlinear neural dynamics. Rather than predefining representational domains (e.g., power, phase, or connectivity), HeteroRC embeds neural time series signals into a recurrent state space with fixed random connectivity and heterogeneous intrinsic time constants. The use of heterogeneous intrinsic time constants is motivated by a growing body of empirical and theoretical work demonstrating that neural dynamics in the cortex unfold across multiple, partially overlapping timescales. Single-neuron recordings have revealed substantial variability in intrinsic and effective time constants, supporting the idea that cortical circuits act as a reservoir of temporal integration windows capable of maintaining information over different durations [50]. At the population and systems level, recent work has shown that intrinsic timescales are organised hierarchically across cortical areas, with progressively longer integration timescales in higher-order regions, facilitating reliable signal propagation and accumulation of task-relevant information [47, 49]. Theoretical analyses further suggest that heterogeneity in neuronal time constants is not a nuisance but a computational resource that shapes population dynamics and expands the repertoire of representational transformations available to neural circuits [48]. By incorporating a distribution of intrinsic time constants, HeteroRC introduces a biologically motivated inductive bias that reflects multiscale cortical dynamics, allowing information distributed over fast and slow temporal regimes to be accessed by linear decoding without assuming instantaneous signal expression.

A known limitation of recurrent decoding approaches is that temporal integration of past inputs can introduce systematic temporal smearing, reducing the precision with which the timing of informative neural events can be recovered [42]. To mitigate this effect in offline analyses, HeteroRC incorporates bidirectional temporal processing, in which neural time series are processed both forward and backward in time, and the resulting reservoir states are combined to cancel direction-specific temporal lags. We explicitly note that this bidirectional, multiplicative fusion is a non-causal technical strategy rather than a biologically plausible mechanism. Conceptually analogous to zero-phase filtering in standard signal processing [56], it is employed here to enhance temporal resolution during offline decoding while retaining the expressive benefits of recurrent integration. For applications requiring strict biological plausibility or real-time, online decoding (e.g., brain–computer interfaces), the framework readily defaults to a causal, unidirectional approach. Ultimately, by training only a regularised linear readout, the framework balances expressive power with data efficiency and interpretability [18], making it well suited to the small-sample regimes and inferential goals typical of cognitive neuroscience experiments.

The decoding results point to a common principle: task-relevant information in neural time-series is not uniformly expressed as transient, phase-locked responses, but often emerges through sustained, trial-variable neural dynamics. Linear classifiers applied to sensor-level amplitude traces are well suited to decoding evoked responses, as evidenced by both simulations and empirical data, where linear decoding captured early, stimulus-locked information with high temporal precision [14, 16, 17]. However, beyond this initial evoked regime, decoding performance with linear models rapidly declined, suggesting limited sensitivity to neural information distributed over time or expressed in non-phase-locked dynamics. In contrast, HeteroRC consistently recovered task-relevant information across a broader range of neural regimes. In the motor imagery dataset, HeteroRC maintained robust decoding throughout extended imagery periods in the absence of external stimulation, consistent with the view that internally generated motor representations are supported by sustained and induced neural dynamics rather than brief cue-locked responses [7, 19, 57]. Similarly, in the attentional priority mapping dataset, HeteroRC decoded spatial priority information both following a visual impulse and during no-ping intervals, indicating that learned priority representations remained accessible even when they were not expressed as overt phase-locked responses. These convergent findings across tasks suggest that many neural representations previously considered difficult to decode may persist as continuous, dynamically evolving states that are poorly matched to the assumptions of instantaneous amplitude-based linear decoding. Together, the simulation and empirical results indicate that the apparent boundaries of decodability in neural time-series signals are shaped not only by the presence or absence of neural information, but by how that information is dynamically expressed and sampled by the decoding model. By integrating neural activity over time while preserving temporal precision, HeteroRC provides access to representational formats that extend beyond evoked responses and are central to internally generated and learned cognitive states.

Beyond decoding performance, an important contribution of the present work lies in the interpretability framework that links successful decoding to identifiable neural signal components. In the context of EEG/MEG/LFPs decoding, improved accuracy alone provides limited insight into how information is represented in neural activity. By projecting decoding weights back into reservoir space and further mapping informative reservoir dynamics to latent virtual sources and sensor-level patterns, the present framework constrains decoding results to be interpretable in terms of temporal, spectral, and spatial neural signatures. Applying this framework revealed that decoding success in different task contexts relied on distinct classes of neural dynamics. In the motor imagery dataset, informative components reflected a combination of sustained temporal dynamics, oscillatory power modulations in sensorimotor rhythms, and aperiodic spectral changes. In the attentional priority mapping dataset, interpretation distinguished between evoked, phase-locked responses following an external impulse and induced, non-phase-locked dynamics supporting decoding in the absence of perturbation. The fact that Ping and No-Ping decoding relied on partially distinct neural signal components provides convergent evidence that HeteroRC accesses multiple representational regimes rather than exploiting a single dominant feature. By extending this interpretation framework to the group level via sensor-space spatial matching, we further demonstrated that these distinct mechanistic motifs are conserved across the cohort, reflecting a generalised neurophysiological phenomenon rather than an idiosyncratic, single-subject artifact. More broadly, these results illustrate how interpretability can serve as a critical safeguard for decoding-based inference in neuroscience. By revealing which neural dynamics support decoding under different conditions, both at the individual and group levels, the interpretation framework enables researchers to assess whether decoding results are consistent with known physiological mechanisms and task demands. In this way, interpretability is not merely a descriptive add-on, but an essential component for drawing meaningful conclusions about neural representations from time-resolved decoding analyses.

While recent advances in neural decoding have increasingly explored large-scale nonlinear models, including CNNs, RNNs, and transformer-based architectures, these approaches are typically optimised for epoch-wise classification and often obscure fine-grained temporal correspondence through hierarchical convolution and pooling operations, complicating analyses of representational dynamics and cross-temporal generalisation [40, 55, 58]. Moreover, their effectiveness commonly depends on large training datasets and extensive hyperparameter optimisation, which are frequently incompatible with the small-sample regimes, inter-subject variability, and interpretive goals characteristic of neuroscience experiments [40, 41]. Our comparative simulations directly substantiate these concerns. Specifically, we showed that standard sequence-to-sequence ANN architectures are highly susceptible to pronounced temporal lags. Because recurrent networks (such as LSTMs) continuously accumulate all preceding information, and unmasked Transformers globally integrate both past and future context, their resulting decoding time courses inherently misalign with the ground-truth temporal windows of the underlying neural signals. Although restricting the temporal integration horizon via a sliding-window approach successfully optimises the temporal precision of these trainable models, their overall decoding accuracy and temporal sharpness still underperform relative to HeteroRC. In comparison to these models, HeteroRC occupies a complementary position in the modelling landscape. By relying on fixed random recurrent connectivity and training only a linear readout, HeteroRC avoids large-scale parameter optimisation while still enabling rich temporal representations through recurrent dynamics. This design yields a favourable trade-off between representational richness, data efficiency, and interpretability, and substantially reduces computational and memory demands relative to fully trainable deep learning models [42, 44]. Rather than approximating an optimal end-to-end decoder, HeteroRC provides a principled and tractable state-space transformation aligned with known properties of neural dynamics, supporting time-resolved and hypothesis-driven analyses in low-data, high-variability settings typical of electrophysiological research.

In conclusion, the HeteroRC framework provides an interpretable and computationally efficient foundation for time-resolved neural decoding. Its lightweight and interpretable architecture, together with its compatibility with time-resolved decoding, makes it well-suited for a broad range of electrophysiological applications, including but not limited to studies of internally generated cognitive states and latent neural representations. To facilitate reproducibility and adoption by the community, the full HeteroRC decoding and interpretation codes are made openly available as a documented, open-source software package on GitHub (https://github.com/rl671/heterorc). We anticipate that this resource will enable systematic investigation of how neural information is dynamically represented over time and promote methodological advances in neural decoding beyond phase-locked responses, with relevance for both neuroscience and brain–computer interface research.

## Methods

### Simulated datasets

To systematically evaluate decoding sensitivity to distinct classes of neural dynamics, we generated synthetic datasets mimicking commonly studied electrophysiological regimes, including stimulus-locked evoked responses, non-phase-locked oscillatory activity, ISPC, and aperiodic (1/*f*) activity. Simulations were designed to isolate each regime while maintaining realistic signal-to-noise characteristics and spatial structure, allowing controlled comparisons between decoding models. All codes used in this study can be found on GitHub (https://github.com/rl671/heterorc).

Each simulated dataset consisted of 30 independent “subjects”. For each subject, we generated two-class classification data with 40 trials per class (80 trials total). Signals were simulated for a standard 32-channel EEG montage, sampled at 100 Hz, over a time window from 0 to 800 ms. Channel labels and regional groupings followed the international 10–20 EEG system. Frontal channels were defined as electrodes with labels beginning with “F,” whereas posterior channels were defined as electrodes with labels beginning with “P,” “O,” or “CP.” For simulations of evoked responses, induced oscillatory activity, and aperiodic spectral modulations, task-related signals were injected exclusively into posterior channels, with frontal channels serving as controls. In the ISPC condition, oscillatory signals were injected into both frontal and posterior channel groups, and class differences were implemented by modulating the strength of phase synchronisation between these regions.

All simulations incorporated realistic background noise composed of a mixture of spatially uncorrelated white noise and temporally correlated pink (1/*f*) noise. For each trial and channel, white Gaussian noise and pink noise were combined with fixed weights (0.4 and 0.6, respectively) and scaled to yield unit-variance signals prior to task-related modulation. This background noise was present in all conditions and was identical across classes, ensuring that classification performance depended exclusively on the injected physiological signal of interest.

Five signal regimes were simulated independently. Across all regimes, task-related modulations were confined to a post-stimulus time window between approximately 200 and 600 ms, with modest trial-to-trial temporal jitter to avoid perfectly aligned onsets and offsets. Across all simulated regimes, class identity was encoded via a moderate difference in the target feature (e.g., amplitude, phase synchronisation, or aperiodic parameters) embedded in a physiological noise background comprising a mixture of 1/f pink and white noise. The resulting macroscopic signal differences are shown in the univariate contrasts (Figure 2).

To simulate stimulus-locked evoked activity, class-discriminative signals were implemented as Gaussian-shaped amplitude deflections time-locked to stimulus onset. The evoked response peaked around 400 ms post-stimulus, with small trial-to-trial temporal (± ∼25 ms) and amplitude jitter. Although this temporal jitter was introduced to approximate physiological variance, the macroscopic amplitude deflections remained largely consistent across trials, ensuring that discriminative information was predominantly phase-locked to the event.

Induced activity was simulated as transient oscillatory bursts with a Gaussian temporal envelope (duration ≈ 400 ms) at a target frequency (default 10 Hz). Oscillatory phase was randomised independently on each trial, rendering the signal non-phase-locked at the sensor level. Class information was encoded solely in oscillatory amplitude. Bursts had modest trial-wise variability in frequency (±0.5 Hz) and amplitude to approximate physiological variability. To assess frequency generality, additional simulations were performed at lower and higher carrier frequencies (5, 15, and 25 Hz), while all other parameters were held constant.

To model phase-based functional connectivity (i.e., ISPC), oscillatory bursts were simultaneously injected into frontal and posterior channel groups. For one class, the phase difference between frontal and posterior signals was tightly clustered (high phase synchronisation), whereas for the other class phase differences were broadly distributed. Importantly, marginal oscillatory power was matched across classes; discriminative information was carried exclusively by inter-regional phase consistency rather than local amplitude or evoked responses. As for induced activity, ISPC simulations were primarily conducted at 10 Hz, with additional carrier frequencies (5, 15, and 25 Hz) examined in supplementary analyses to verify frequency-independent decoding performance.

Aperiodic neural activity was simulated using a physiologically motivated spectral rotation model. For each channel, broadband noise was filtered to produce 1/f-like spectra with controllable slope (exponent) and intercept (offset in log-power space). Class differences were implemented as changes in either the spectral slope or intercept, pivoting around a fixed frequency (default 10 Hz). Aperiodic modulations were applied using a smooth temporal mask. Outside this interval, all channels exhibited identical baseline aperiodic structure.

### Motor imagery dataset

We evaluated HeteroRC on the publicly available Graz motor imagery dataset from BCI Competition IV (2008), data set 2a (https://www.bbci.de/competition/iv/#datasets) [51]. The dataset comprises EEG recordings from nine healthy participants performing a cue-based motor imagery task. Each participant completed two recording sessions on separate days; one session was designated as training data with class labels provided, and the other as evaluation data.

Participants performed four types of motor imagery: left-hand movement, right-hand movement, both feet movement, and tongue movement. Each session consisted of six runs, and each run included 48 trials (12 trials per class), yielding 288 trials per session. At the beginning of each trial, a fixation cross appeared on the screen accompanied by a brief auditory warning tone. After 2 s of fixation, a visual cue in the form of an arrow pointing either to the left, right, down, or up indicated the motor imagery class and remained on the screen for 1.25 s. Participants were instructed to continue the motor imagery task for 2.75 s until the fixation cross disappeared. No feedback was provided during the task.

EEG was recorded using 22 Ag/AgCl electrodes arranged according to the international 10–20 system, with inter-electrode distances of approximately 3.5 cm. Signals were recorded monopolarly with the left mastoid as reference and the right mastoid as ground. Data were sampled at 250 Hz and bandpass-filtered online between 0.5 and 100 Hz, with an additional 50 Hz notch filter applied to suppress line noise. Three additional EOG channels were recorded for artefact monitoring but were not used for decoding analyses.

Offline preprocessing was intentionally kept minimal. EEG data were bandpass-filtered between 0.5 and 30 Hz, downsampled to 100 Hz, and re-referenced to the common average. No additional artefact correction procedures were applied.

### Attentional priority mapping dataset

We further evaluated HeteroRC on a publicly available EEG dataset investigating history-driven attentional priority maps (https://osf.io/v7yhc), originally reported by Duncan, van Moorselaar [36]. In this study, 24 participants performed a visual search task designed to induce statistical learning of spatial target probabilities, leading to the implicit formation of an attentional priority map. One participant was excluded due to a technical failure during data loading, resulting in a final sample of 23 participants included in the analyses.

Participants completed a variant of the additional singleton task in which targets appeared more frequently at one of four cardinal locations (up, down, left, or right) across blocks, while other locations were less probable. Critically, on half of the trials, a task-irrelevant but salient visual “ping” stimulus was presented during the intertrial interval, whereas no such stimulus was shown on the remaining trials. This design explicitly allows comparison of ping and no-ping conditions while holding task demands constant.

EEG was recorded from 64 scalp electrodes arranged according to the international 10–10 system using a BioSemi ActiveTwo system. In the original study, EEG signals were re-referenced to the common average, high-pass filtered at 0.01 Hz. Epochs containing pronounced EMG or muscle-related artefacts were then identified and removed. Independent component analysis (ICA) was subsequently applied to remove ocular and other stereotypical artefacts, and trials containing eye movements or saccades were further excluded based on concurrent eye-tracking data.

To ensure consistency across datasets and decoding analyses, we applied additional preprocessing steps to the provided data. Specifically, EEG signals were bandpass-filtered between 0.5 and 30 Hz and downsampled to 100 Hz. No further artefact rejection or channel selection was performed beyond the preprocessing implemented in the original dataset (see Duncan, van Moorselaar [36] for details). No time-domain baseline correction was applied in the decoding analyses, in order to preserve neural activity preceding the ping stimulus.

### Univariate analyses of simulated data

To characterise the signal properties of the simulated datasets independently of decoding, we performed standard univariate analyses at the subject level and summarised results across simulated subjects. For each simulation mode, data were generated for multiple simulated subjects with identical generative parameters but independent noise realisations.

Evoked responses were computed by averaging epoched sensor-level signals across trials and across all electrodes within each subject and condition, yielding a global mean time course. Power spectral density (PSD) was estimated using Welch’s method over the entire time window and averaged across trials and electrodes within each subject, with power expressed in decibels. To quantify phase-based interactions in the simulated data, we computed ISPC, also referred to as the phase-locking value (PLV), between predefined frontal and posterior electrode groups. ISPC was computed as the magnitude of the average unit phase-difference vector across trials within a predefined temporal window (0.2–0.6 s):

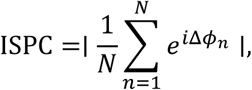

where Δ*ϕ*_*n*_ denotes the phase difference between frontal and posterior regions for trial *n*, evaluated at the midpoint of the temporal window, and *N* is the number of trials. ISPC values range from 0 (random phase differences across trials) to 1 (perfect phase alignment). In addition, per-trial phase differences between regions were pooled across subjects to visualise the distribution of phase offsets. For all univariate measures, subject-level results were averaged across simulated subjects, and variability was quantified using the standard error of the mean.

### Time-resolved decoding and cross-temporal generalisation using LDA and SVM

As baseline decoding approaches, we employed LDA and linear SVM, which are widely used in time-resolved EEG/MEG/LFPs decoding. To evaluate instantaneous signal decodability, decoding was performed independently at each discrete time point using a sliding-estimator approach, such that classifiers were trained and evaluated on sensor-level amplitude patterns across all sensors at each individual time sample. Prior to classification, features were standardised using z-scoring based on the training data only. For decoding based on conventional time-frequency features (Figure S2), we extracted alpha-band power (8–12 Hz) using two classic approaches. In the first, we computed spectral power via a Morlet wavelet transform (with cycles dynamically set to *f*/2), averaging the resulting estimates across 8–12 Hz. In the second, we applied an 8–12 Hz bandpass filter to the raw signals and extracted instantaneous power via a Hilbert transform (calculated as the squared absolute value of the analytical signal). For both approaches, the resulting power time series were standardised and submitted to the identical sliding-estimator LDA pipeline.

For simulated datasets and the attentional priority mapping dataset, time-resolved decoding accuracy was estimated using stratified 5-fold cross-validation. At each time point, classifiers were trained on the training folds and evaluated on held-out data, yielding a decoding accuracy time course that was then averaged across folds.

For the motor imagery dataset, decoding performance was evaluated using a fixed train–test split defined by recording sessions. Classifiers were trained on data from the training session and tested on data from a separate recording session collected on a different day, providing a stringent assessment of cross-session generalisation without cross-validation.

To characterise the temporal stability and transformation of neural representations, we performed cross-temporal generalisation analyses for both linear decoders for the motor imagery dataset. In this approach, classifiers were trained at a given time point and tested at all other time points, yielding a two-dimensional matrix of decoding accuracy as a function of training time and testing time. Cross-temporal generalisation was evaluated using the same cross-session train-test split as time-resolved decoding, with classifiers trained on one session and tested on the other. This analysis assesses the temporal generalisation of neural representations across time under strict cross-session generalisation constraints.

### Heterogeneous Reservoir Computing (HeteroRC)

#### Overview

HeteroRC is a temporal decoding framework based on an echo state network for extracting task-relevant information from multichannel electrophysiological time series. The model projects neural signals into a high-dimensional recurrent state space with heterogeneous intrinsic timescales, followed by a linear readout trained to decode experimental variables. By combining recurrent dynamics with explicit temporal heterogeneity, HeteroRC is sensitive to both stimulus-locked, phase-consistent activity and non-phase-locked neural dynamics that unfold over longer or variable temporal scales.

HeteroRC differs from conventional recurrent neural networks in three key respects: (i) both the input-to-reservoir weights and the recurrent weights within the reservoir are fixed and not optimised during training, (ii) non-linear transformations arise solely from the intrinsic reservoir dynamics rather than from learned representations, and (iii) temporal heterogeneity is explicitly imposed by assigning distinct intrinsic time constants to reservoir units. As a result, HeteroRC functions as a structured dimension expansion of the input time series, enabling information carried by evoked responses as well as induced, non-phase-locked dynamics to be represented within a common state space and decoded using a linear readout.

#### Input representation and normalisation

Let **X** ∈ ℝ^*N*×*C*×*T*^denote the epoched neural time-series data, where *N* is the number of trials, *C* is the number of channels, and *T* is the number of time points. For each trial *n*, the multivariate time series **x**_*n*_(**t**) ∈ ℝ^*C*^ is provided directly to the reservoir at each time point **t**. To ensure strict separation between training and testing data, input normalisation (across all trials, channels, and time points) was performed independently within each cross-validation fold. Specifically, input amplitudes were scaled using the 99th percentile of the absolute signal amplitude computed from the training data only, and the same global scaling factor was applied to the corresponding test data. This normalisation constrains input magnitudes to a consistent dynamic range, ensuring that reservoir units operate within the sensitive (non-saturated) regime of the tanh nonlinearity.

#### Reservoir architecture and dynamics

The reservoir consists of *R* recurrent units with leaky-integrator dynamics. The state of the reservoir at time **t** for trial *n* is denoted by **r**_*n*_(**t**) ∈ ℝ^*R*^. Reservoir dynamics evolve according to the discrete-time update equation:

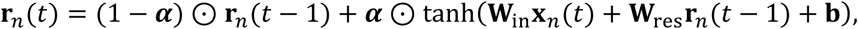

where **W**_in_ ∈ ℝ^*R*×*C*^ is a fixed random input projection matrix, **W**_res_ ∈ ℝ^*R*×*R*^ is a fixed sparse recurrent weight matrix (10% connectivity density), and **b** ∈ ℝ^*R*^ is a bias vector. The operator ⊙ denotes element-wise multiplication, and **α** ∈ ℝ^*R*^ is a vector of unit-specific leak rates. The input weights **W**_in_, recurrent weights **W**_res_, and biases **b** are randomly initialized and remain fixed throughout training and testing. Recurrent weights are scaled to achieve a target spectral radius less than unity, ensuring echo-state stability.

#### Heterogeneous intrinsic timescales

A defining feature of HeteroRC is the explicit introduction of heterogeneous intrinsic timescales across reservoir units. Each reservoir unit *i* is assigned a time constant *τ*_*i*_, drawn from a log-normal distribution and constrained to lie within a physiologically plausible range [*τ*_min_, *τ*_max_]. Distribution parameters are chosen such that the mode of the distribution corresponds to a biologically motivated timescale (e.g., *τ*_mode_ = 10 ms), while allowing a heavy tail that captures slower dynamics. The leak rate for each unit is derived from its time constant according to:

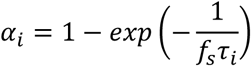

where *f*_*s*_ is the sampling frequency. Under this formulation, each reservoir unit effectively acts as a low-pass temporal filter with a cutoff frequency *f*_*c*_ = (2*πτ*_*i*_)^−1^. The reservoir therefore implements a multiscale temporal filter bank, enabling integration of input information over diverse temporal windows without explicit frequency-domain decomposition.

#### Bidirectional temporal processing

To enhance sensitivity to temporally extended and temporally symmetric patterns, HeteroRC employs bidirectional temporal processing. Each trial is processed independently in both forward and backward temporal directions using identical reservoir parameters. Backward reservoir states are obtained by applying the reservoir to time-reversed input sequences. Forward and backward reservoir states are combined at each time point using element-wise multiplication:

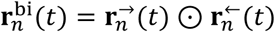

This multiplicative fusion emphasises temporally consistent structure across directions while suppressing transient, direction-specific noise, and preserves the fixed-weight nature of the reservoir. Alternative processing strategies, including unidirectional reservoirs and bidirectional fusion via element-wise averaging, were also evaluated in supplementary analyses (Figure S3). Unless otherwise stated, all main analyses in the text use bidirectional processing with multiplicative fusion.

#### Linear readout and time-resolved decoding

Decoding is performed using a linear classifier trained on the bidirectional reservoir states at each time point. Let 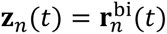 denote the final reservoir representation. Class scores are computed as: 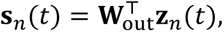, where **W**_out_ is estimated using ridge-regularised linear classification. Time-resolved decoding performance was evaluated using stratified 5-fold cross-validation for simulated datasets and the attentional priority mapping dataset, and using a fixed cross-session train–test split for the motor imagery dataset. Readout weights were trained on the training data and evaluated on held-out data at each time point, yielding a decoding accuracy time course that was averaged across folds or across test samples, respectively. To improve robustness of time-resolved decoding, decision scores were temporally smoothed using a Gaussian kernel (FWHM = 25 ms) prior to class prediction. Cross-temporal generalisation analyses were performed only for the motor imagery dataset and followed the same cross-session train–test split. Readout weights trained at a given time point in the training session were evaluated across all time points in the independent test session, yielding training-by-testing time generalisation matrices. In this decoding procedure, only the readout weights **W**_out_ are trained. All reservoir parameters were fixed and shared between training and testing data. Consequently, decoding performance reflects the expressive capacity of the reservoir dynamics rather than representational learning in the readout.

#### HeteroRC parameter settings

In this study, unless otherwise stated, the following parameter settings were used for all HeteroRC analyses. For simulated datasets, the reservoir size was set to *R* = 350 units. For analyses of real EEG datasets, including the motor imagery and attentional priority mapping datasets, the reservoir size was increased to *R* = 800 units to accommodate the higher dimensionality and variability of empirical data. Across all datasets, the recurrent weight matrix was scaled to a target spectral radius of 0.95. Intrinsic time constants were drawn from a log-normal distribution with mode *τ*_mode_ = 0.01 s and shape parameter *σ* = 0.8, producing a heavy-tailed distribution of timescales. Time constants were constrained to lie within the range [*τ*_min_, *τ*_max_] = [0.002,0.08]s. Bidirectional temporal processing was enabled in all analyses, with forward and backward reservoir states combined using element-wise multiplication. As a general guideline for hyperparameter selection in this framework, *R* should be chosen to be large enough to provide a sufficiently rich high-dimensional space for disentangling complex dynamics. A common heuristic is to set *R* to approximately 10 to 15 times the number of input channels. Additionally, setting the spectral radius just below unity (typically between 0.90 and 0.99) is a typical practice to rigorously enforce the echo state property, ensuring the network maintains a stable, fading memory of past inputs without transitioning into chaotic dynamics.

### Individual-level interpretability framework

To interpret task-relevant information represented within the HeteroRC reservoir, we first developed an individual-level interpretability framework that progressively links reservoir dynamics to sensor-level neurophysiological signals. This framework extends covariance-based activation pattern analysis to recurrent reservoirs and enables separation of heterogeneous temporal, spectral, and spatial contributions to decoding performance.

#### Identification of task-relevant latent dynamics in reservoir space

We first identified reservoir units that contributed most strongly to the decoding decision. In multivariate decoding models, linear readout weights cannot be directly interpreted as feature importance because they reflect both signal encoding and noise suppression induced by feature covariance [18]. To recover task-related activation patterns within the reservoir, we applied a covariance-based transformation of the classifier weights, following the approach introduced by Haufe and colleagues [18] to yield interpretable activation coefficients.

Let **r**(**t**) ∈ ℝ^*R*^ denote the reservoir state vector at time **t**, and 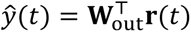 denote the corresponding decision value of the linear readout. Activation patterns **A** were computed as:

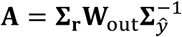

where **∑**_**r**_ is the covariance matrix of reservoir states and **∑***_ŷ_* denotes the covariance of the decision values. This transformation yields activation coefficients reflecting how strongly each reservoir unit encodes task-relevant variance.

For multi-class decoding, a global importance score was assigned to each reservoir unit as the maximum absolute activation coefficient across classes. The dominant class contribution for each unit was defined as the class associated with the largest absolute activation coefficient. To disentangle distinct dynamical mechanisms represented within the high-dimensional reservoir state space, we selected the most informative units and performed functional clustering based on their temporal activity profiles. For individual-level interpretations, we retained the top 25 units by importance. Prior to clustering, the time series of the selected units were z-score standardised to ensure that grouping was driven by the shape of temporal dynamics rather than signal amplitude. We then applied hierarchical clustering using Ward’s method with the Euclidean distance metric. This procedure grouped reservoir units into a small number of functional clusters reflecting distinct temporal and spectral dynamics. For each cluster *k*, a virtual source signal *v*_*k*_(**t**) was constructed as a weighted combination of each constituent reservoir unit *i* within cluster *C*_*k*_:

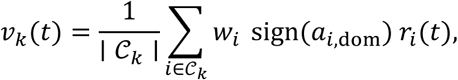

where *w*_*i*_ denotes the unit importance score and the sign term enforces consistency with the dominant class contribution. These virtual sources represent low-dimensional summaries of task-relevant reservoir dynamics.

For each virtual source *v*_*k*_(**t**), we quantified its task-relevant dynamics in the time, time–frequency, and spectral domains. First, event-related responses were computed by averaging *v*_*k*_(**t**) within each experimental condition across trials, yielding class-wise virtual-source evoked responses. Second, time–frequency representations were estimated using complex Morlet wavelets over frequencies from 2 to 40 Hz (2 Hz steps). The number of cycles scaled with frequency (*n*_cycles_ = *f*/2), and power was computed for each condition and then averaged across trials. To facilitate interpretation of induced changes, power was baseline-corrected in decibel units by referencing each frequency bin to the mean power in a pre-stimulus baseline window (i.e., 10log _10_(*P*/*P*_base_)). Third, PSD was computed from *v*_*k*_(**t**) using Welch’s method and condition-specific PSDs were averaged across trials and visualised on a log-frequency axis. To dissociate aperiodic and periodic components, spectra were parameterised using FOOOF [28] over 2–40 Hz with a maximum of three oscillatory peaks, and the fixed aperiodic mode was used to summarise 1/f-like structure.

#### Back-projection of reservoir dynamics to sensor space

In this stage, we linked task-relevant reservoir dynamics back to the sensor space to recover their spatial origins. Under the linear forward model of volume conduction, the spatial topography of a latent source can be estimated by its covariance with sensor-level measurements [18, 59, 60]. To capture the spatial expression of the reservoir dynamics specifically during the decision-making process, we restricted our analysis to a focused time window (±100 ms) centred around the peak decoding accuracy. We estimated a spatial projection **p**_*k*_ ∈ ℝ^*C*^ for each virtual source *v*_*k*_(**t**) by computing its covariance with the raw sensor signals **x**(**t**) within this window:

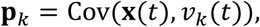

The resulting spatial patterns **p**_*k*_ represent the topographical distribution of the specific latent dynamics (i.e., the virtual source signal) corresponding to each functional cluster.

### Group-level interpretability framework

To determine whether the latent representational dynamics observed at the single-subject level were conserved across the cohort, we extended the interpretability framework to perform group-level inference. Because randomly initialised reservoirs yield specific and incommensurable latent spaces, direct cross-subject alignment of reservoir units is challenging. To overcome this, we developed a sensor-space spatial matching approach, leveraging the fact that the physical EEG sensor array is anatomically standardised across participants. For each participant, we extracted the 10 most informative reservoir units and computed their spatial topographies (sensor-level covariance maps) within a ±100ms window centred on the participant-specific peak decoding time. As two reservoir units may capture identical neural dynamics but with inverted polarities, we applied a spatial sign-alignment procedure: for each unit’s topography, we identified the sensor with the maximum absolute covariance and multiplied the entire topographical map by -1 if this peak value was negative. The corresponding sign-flip coefficient was retained for subsequent signal reconstruction. These sign-aligned maps were then z-scored to ensure that matching was driven strictly by the underlying spatial distribution (i.e., cortical generator patterns) rather than absolute covariance amplitude. Standardised spatial maps from all selected units across all participants were concatenated into a single global feature matrix. We then applied hierarchical clustering (Ward’s method, Euclidean distance) to group the units into distinct global spatial clusters.

To reconstruct the group-level neurophysiological dynamics, we mapped the globally clustered units back to their respective participants. Participant-specific virtual sources were reconstructed by computing a weighted sum of the constituent unit time series. During this reconstruction, each unit’s time series was multiplied by its previously determined sign-flip coefficient. Subject-level temporal, spectral, and time–frequency representations were then extracted from these aligned virtual sources and averaged across participants to yield grand-average profiles. Finally, to generate the grand-average spatial topography for each global cluster, the raw spatial covariance maps of the aligned virtual sources were averaged across participants and z-scored.

### Time-resolved decoding using RNN, LSTM, transformer, and EEGNet

To compare HeteroRC against fully trainable ANNs, we implemented time-resolved decoding pipelines based on RNNs, LSTM networks, Transformers, and EEGNet. We restricted this comparison to the simulated phase-locked evoked responses because these highly consistent signals represent the most straightforward scenario for ANNs to achieve stable performance. The data generation and cross-validation procedures (stratified 5-fold) were identical to those described above. To preserve strict train/test separation, input amplitudes were normalised independently within each fold using the 99th percentile of the absolute amplitude computed exclusively on the training data, matching the HeteroRC preprocessing pipeline.

For the RNN, LSTM, and Transformer, we evaluated two decoding formulations. In the standard (sequence-to-sequence) formulation, the full trial sequence was provided as input, and the models were trained to produce a continuous trajectory of class predictions, generating a discrete decision at every individual time step based on the accumulated signal history. In the windowed (sequence-to-one) formulation, decoding was performed using sliding windows of 10 samples (100 ms). To enable evaluation at early time points, inputs were zero-padded at the epoch onset. At each time step, windowed models output a single prediction based on the preceding 100 ms context. EEGNet was evaluated exclusively using this windowed formulation to respect its convolutional architecture.

Regarding model architectures, the standard RNN and LSTM models consisted of a single recurrent layer (with *tanh* nonlinearity for the RNN) followed by dropout and a linear classification head. The standard Transformer projected sensor-level inputs to a latent embedding, applied sinusoidal positional encoding, and processed the sequence through a two-layer Transformer encoder prior to linear classification. The windowed variants retained these core architectures but aggregated temporal information to produce a single prediction per window: the windowed RNN and LSTM utilised the final hidden state, while the windowed Transformer applied mean pooling across the encoded time dimension. Hidden dimensionality was fixed at 32 units across all RNN, LSTM, and Transformer models to match representational capacity.

All models were implemented in PyTorch and trained on a NVIDIA GeForce RTX 4090 GPU. The RNN, LSTM, and Transformer models were trained using the Adam optimiser (learning rate = 0.002, weight decay = 0.0001) for 30 training iterations with a dropout rate of 0.5. The batch size was set to 64 for standard models and 128 for windowed models. EEGNet was trained using the AdamW optimiser (learning rate = 0.005, weight decay = 0.001) for 25 training iterations with a batch size of 64. Time-resolved decoding accuracy was computed independently at each time point by comparing predicted and ground-truth class labels, yielding a time-resolved accuracy profile mirroring the evaluation approach used for HeteroRC.

### Statistical analysis

Statistical significance of time-resolved decoding accuracy and cross-temporal generalisation matrices was assessed using nonparametric cluster-based permutation tests across subjects, as implemented for one-sample tests against chance level. This approach controls for multiple comparisons across time points while making minimal assumptions about the underlying distributions.

Comparisons of peak decoding accuracy across decoding methods were performed using paired-sample *t*-tests (two-tailed), applied separately to predefined temporal windows and conditions. Unless otherwise stated, statistical tests were conducted at a significance threshold of *p* < 0.05.

## Acknowledgements

This project was supported by UKRI MRC intramural funding MC_UU_00030/15 to A.W. R.L. was supported by a Gates Cambridge Scholarship (OPP1144) and a postdoctoral fellowship from the Canadian Institutes for Health Research (200883). S.L. was supported by Vetenskapsrådet under award 2023-00493. For the purpose of open access, the author has applied a Creative Commons Attribution (CC BY) licence to any Author Accepted Manuscript version arising from this submission.

## Author contributions

**R.L.**: Conceptualisation, Methodology, Software, Formal analysis, Data Curation, Writing - Original Draft, Writing - Review & Editing, Visualisation, Project administration. **S.L.**: Software, Validation, Formal analysis, Writing - Review & Editing. **Y. L.**: Software, Writing - Review & Editing. **J.D.**: Writing - Review & Editing. **R.N.H.**: Methodology, Writing - Review & Editing. **A.W.**: Methodology, Writing - Review & Editing, Supervision, Funding acquisition.

## Supplementary

**Figure S1.**
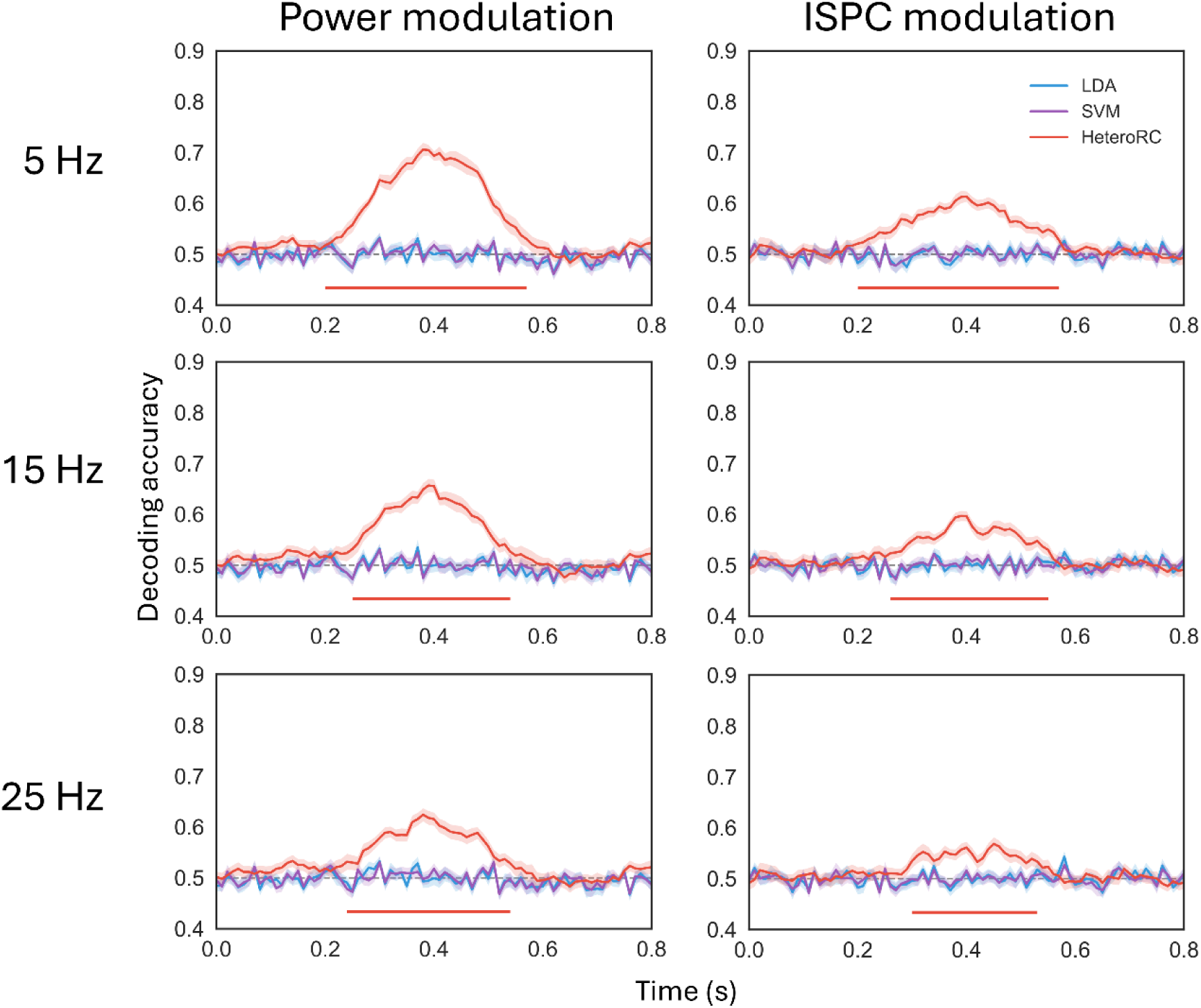
HeteroRC decodes non-phase-locked neural dynamics across a broad range of frequencies. Time-resolved decoding accuracy for simulated induced oscillatory power (left) and inter-site phase clustering (ISPC; right) when task-relevant modulations were centred at frequencies (5 Hz, 15 Hz, and 25 Hz) different from those used in the main text. HeteroRC robustly decodes task-relevant information across frequencies, during the time window that they were introduced (0.2-0.6s), whereas linear raw amplitude-based decoders remain at chance. This demonstrates that HeteroRC decoding performance does not rely on tuning to a specific oscillatory band. Conventions as Figure 2.

**Figure S2.**
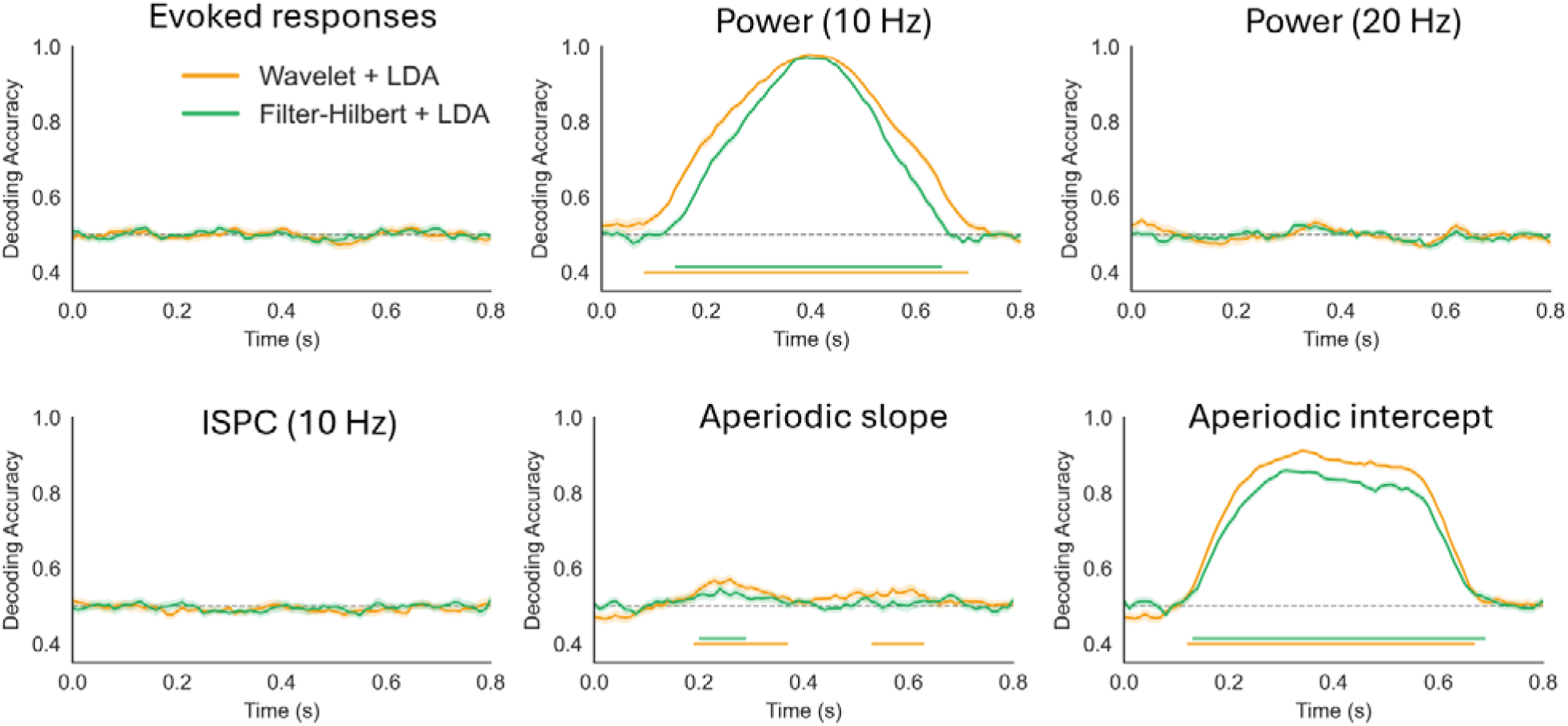
Conventional time-frequency feature extraction restricts decoding sensitivity and induces temporal smearing. Comparison of linear discriminant analysis (LDA) performance applied to *a priori* narrow-band power extracted via Morlet wavelets (8–12 Hz; orange) and a Hilbert filter approach (8–12 Hz; green). Panels display decoding accuracy for simulated data in which the two classes varied in their phase-locked evoked responses, induced oscillatory power centred at 10 Hz and 20 Hz, inter-site phase clustering (ISPC) at 10 Hz, and aperiodic spectral modulations affecting the 1/f slope and intercept (as in Figure 2). Simulated task-relevant modulations were confined to a 0.2–0.6 s time window. Conventions as in Figure 2.

**Figure S3.**
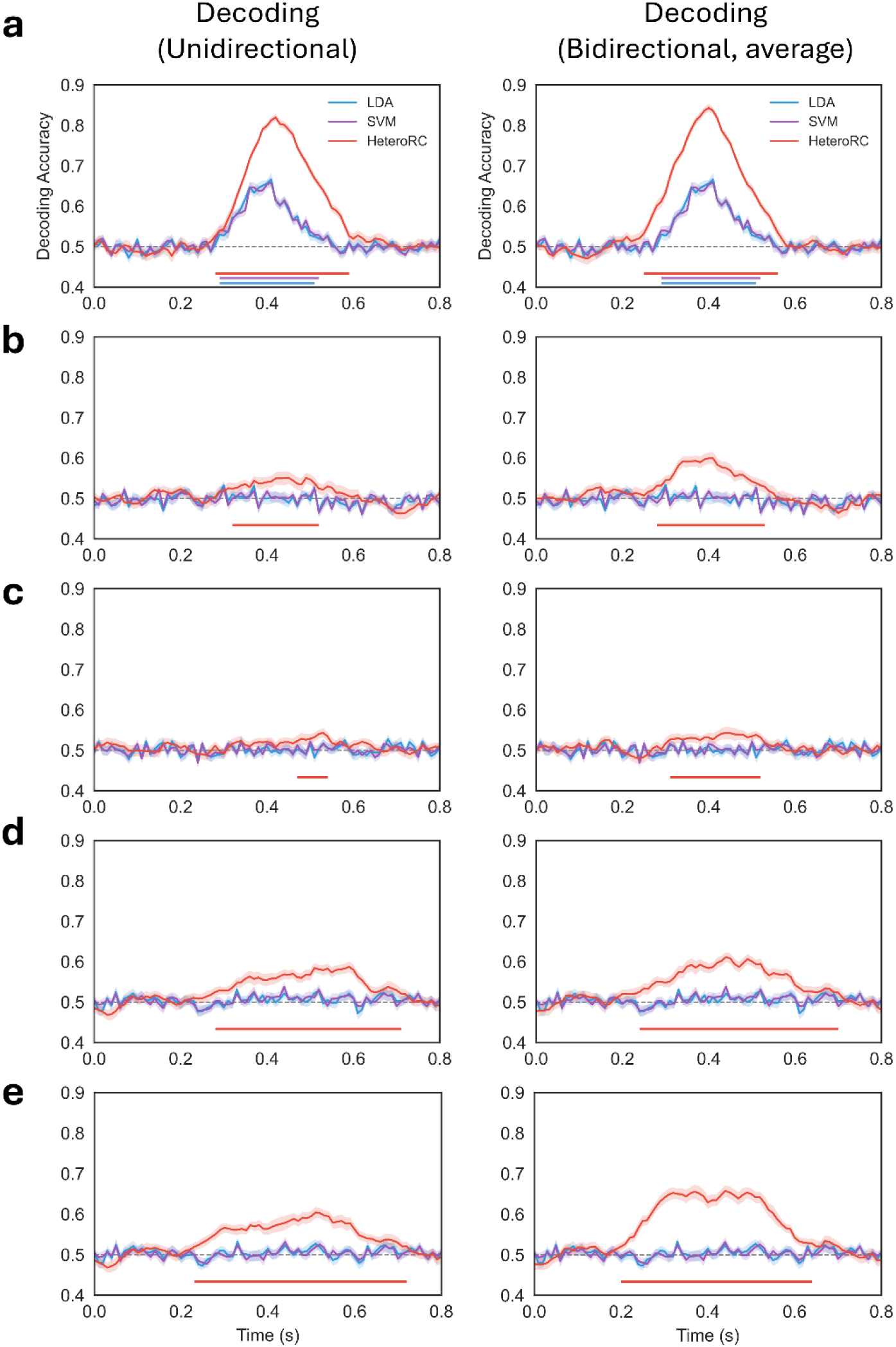
Bidirectional reservoir processing improves temporal precision and reduces decoding smear. Comparison of unidirectional processing (left column) and bidirectional processing (right column) with state averaging for decoding of phase-locked evoked responses (a), induced oscillatory power with randomized phase (b), inter-site phase clustering (ISPC) (c), and aperiodic spectral modulations affecting the 1/f slope (d) and offset (e). Simulated task-relevant modulations were confined to a 0.2–0.6 s time window. Unidirectional processing exhibits systematic temporal lag and smearing in decoding peaks, whereas bidirectional processing improves temporal alignment. Yet, averaging-based fusion shows residual temporal smearing for aperiodic modulations (d and e). In contrast, bidirectional multiplicative fusion (used in the main text, Figure 2) does not exhibit this smearing. Conventions as in Figure 2.

**Figure S4.**
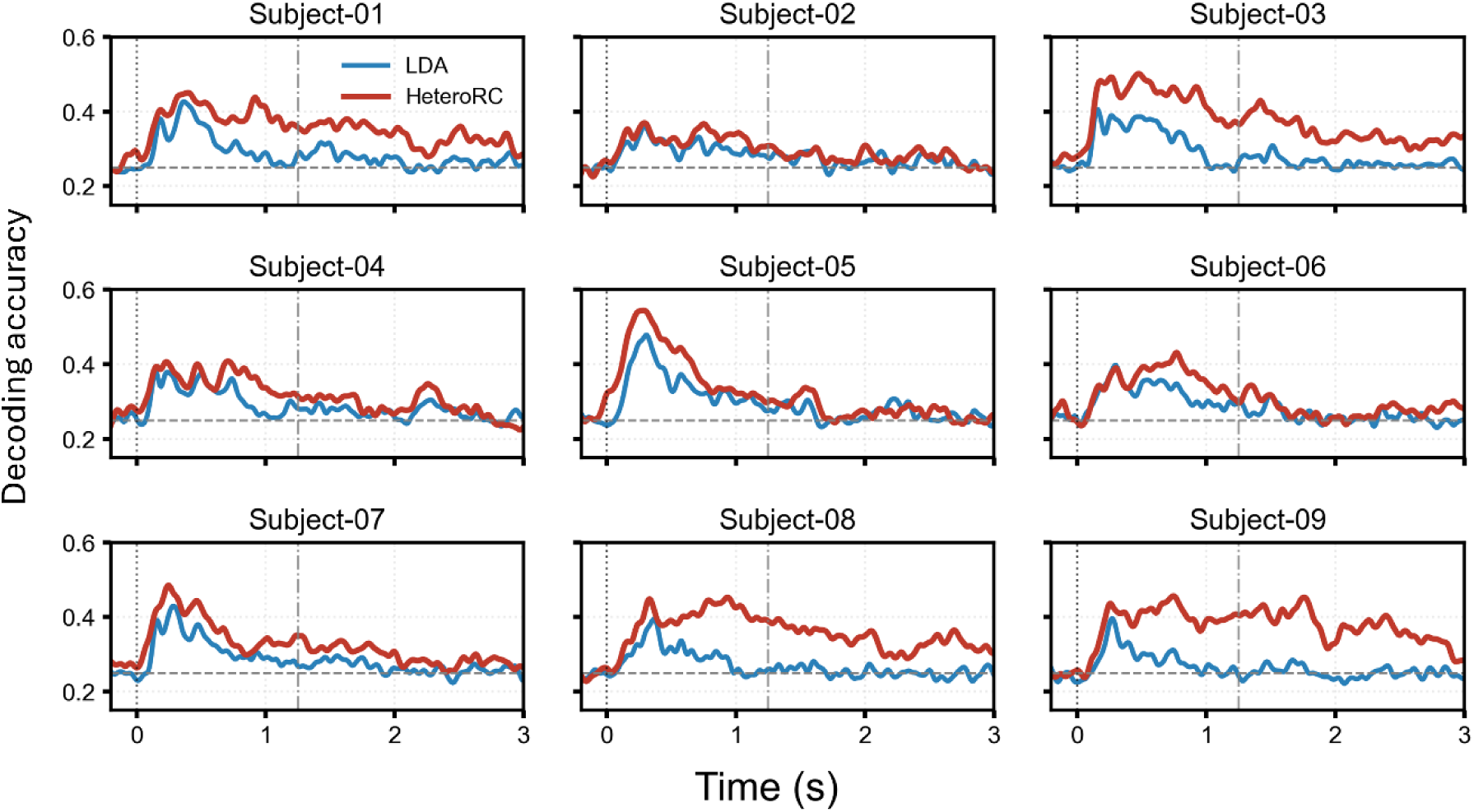
Individual-subject decoding results for the motor imagery dataset. Time-resolved decoding accuracy for each participant in the motor imagery dataset, shown for HeteroRC (red) and linear discriminant analysis (LDA, blue). For visualisation purposes, decoding curves were smoothed with a Gaussian kernel (50 ms FWHM). These plots illustrate the consistency and inter-subject variability of decoding dynamics across participants.

**Figure S5.**
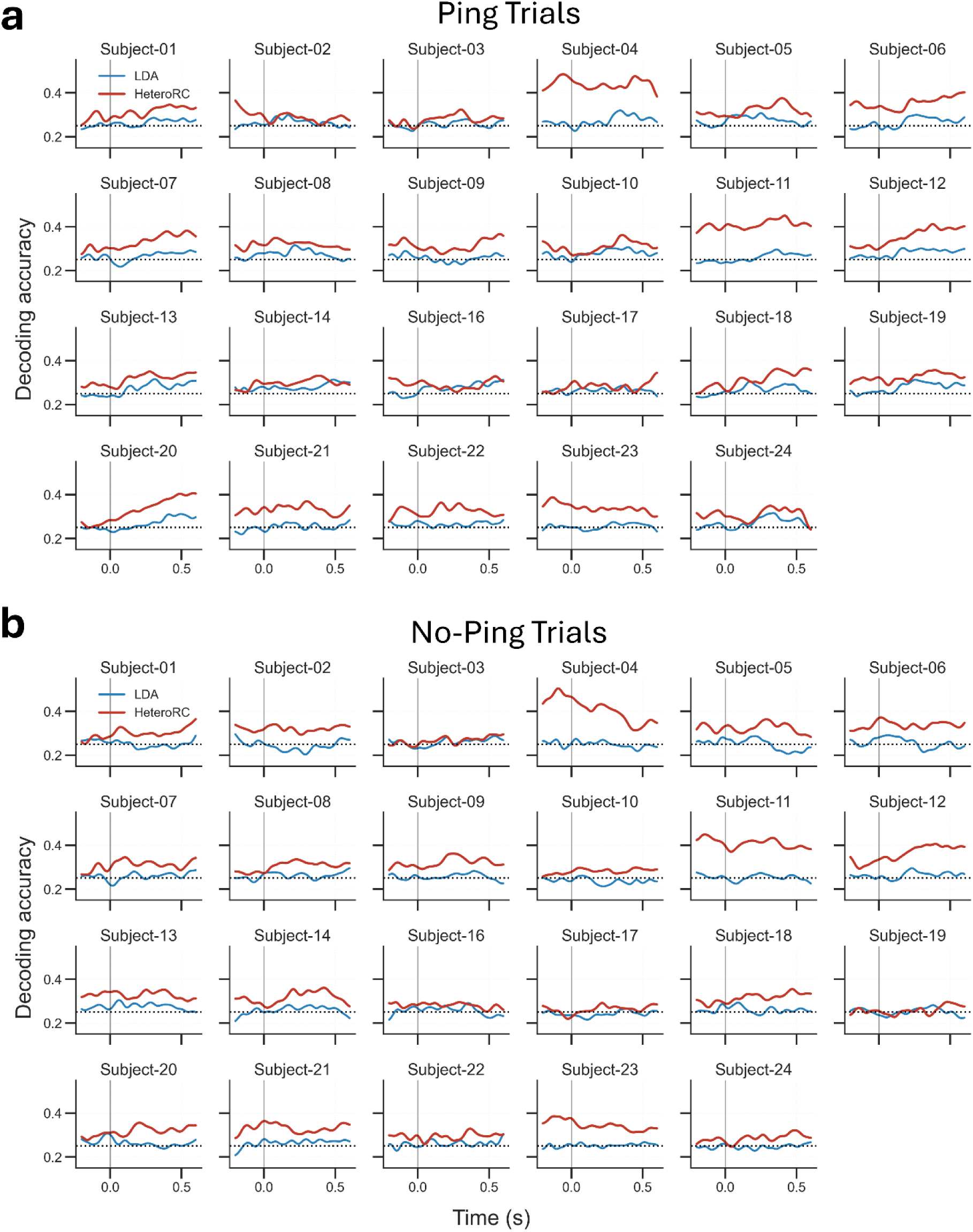
Individual-subject decoding results for the attentional priority mapping task. Time-resolved decoding accuracy for each participant in the attentional priority mapping dataset, shown separately for Ping and No-Ping conditions and for HeteroRC and LDA. Decoding curves were smoothed with a Gaussian kernel (50 ms FWHM) for visualisation.

## References

1. Haxby, J.V., A.C. Connolly, and J.S. Guntupalli, Decoding neural representational spaces using multivariate pattern analysis. Annu Rev Neurosci, 2014. 37: p. 435–56.

2. Norman, K.A., et al., Beyond mind-reading: multi-voxel pattern analysis of fMRI data. Trends Cogn Sci, 2006. 10(9): p. 424–30.

3. Haxby, J.V., et al., Distributed and overlapping representations of faces and objects in ventral temporal cortex Science, 2001. 293(5539): p. 2425–30.

4. Kamitani, Y. and F. Tong, Decoding the visual and subjective contents of the human brain. Nat Neurosci, 2005. 8(5): p. 679–85.

5. Carlson, T.A., P. Schrater, and S. He, Patterns of activity in the categorical representations of objects. J Cogn Neurosci, 2003. 15(5): p. 704–17.

6. Grootswagers, T., S.G. Wardle, and T.A. Carlson, Decoding Dynamic Brain Patterns from Evoked Responses: A Tutorial on Multivariate Pattern Analysis Applied to Time Series Neuroimaging Data. J Cogn Neurosci, 2017. 29(4): p. 677–697.

7. Lotte, F., et al., A review of classification algorithms for EEG-based brain-computer interfaces: a 10 year update. J Neural Eng, 2018. 15(3): p. 031005.

8. Ding, Y., et al., EEG-based brain-computer interface enables real-time robotic hand control at individual finger level. Nat Commun, 2025. 16(1): p. 5401.

9. King, J.R. and S. Dehaene, Characterizing the dynamics of mental representations: the temporal generalization method. Trends Cogn Sci, 2014. 18(4): p. 203–10.

10. Peelen, M.V. and P.E. Downing, Testing cognitive theories with multivariate pattern analysis of neuroimaging data. Nat Hum Behav, 2023. 7(9): p. 1430–1441.

11. Robinson, A.K., G.L. Quek, and T.A. Carlson, Visual Representations: Insights from Neural Decoding. Annu Rev Vis Sci, 2023.

12. Cichy, R.M., D. Pantazis, and A. Oliva, Resolving human object recognition in space and time. Nat Neurosci, 2014. 17(3): p. 455–62.

13. Hebart, M.N. and C.I. Baker, Deconstructing multivariate decoding for the study of brain function. Neuroimage, 2018. 180(Pt A): p. 4–18.

14. Bae, G.Y. and S.J. Luck, Dissociable Decoding of Spatial Attention and Working Memory from EEG Oscillations and Sustained Potentials. J Neurosci, 2018. 38(2): p. 409–422.

15. Lu, R., et al., Parietal alpha stimulation causally enhances attentional information coding in evoked and oscillatory activity. Brain Stimul, 2025. 18: p. 114–27.

16. Renton, A.I., D.R. Painter, and J.B. Mattingley, Optimising the classification of feature-based attention in frequency-tagged electroencephalography data. Scientific Data, 2022. 9(1).

17. Trammel, T., et al., Decoding semantic relatedness and prediction from EEG: A classification method comparison. Neuroimage, 2023. 277: p. 120268.

18. Haufe, S., et al., On the interpretation of weight vectors of linear models in multivariate neuroimaging. Neuroimage, 2014. 87: p. 96–110.

19. Pantazis, D., et al., Decoding the orientation of contrast edges from MEG evoked and induced responses. Neuroimage, 2018. 180(Pt A): p. 267–279.

20. Foster, J.J. and E. Awh, The role of alpha oscillations in spatial attention: limited evidence for a suppression account. Curr Opin Psychol, 2019. 29: p. 34–40.

21. Stecher, R., R.M. Cichy, and D. Kaiser, Decoding the rhythmic representation and communication of visual contents. Trends Neurosci, 2025. 48(3): p. 178–188.

22. Lundqvist, M., et al., Beta: bursts of cognition. Trends Cogn Sci, 2024. 28(7): p. 662–676.

23. Johnson, E.L., et al., A rapid theta network mechanism for flexible information encoding. Nat Commun, 2023. 14(1): p. 2872.

24. Siegel, M., T.H. Donner, and A.K. Engel, Spectral fingerprints of large-scale neuronal interactions. Nat Rev Neurosci, 2012. 13(2): p. 121–34.

25. Fries, P., Rhythms for Cognition: Communication through Coherence. Neuron, 2015. 88(1): p. 220–35.

26. Palva, J.M., et al., Neuronal synchrony reveals working memory networks and predicts individual memory capacity. Proc Natl Acad Sci U S A, 2010. 107(16): p. 7580–5.

27. Albouy, P., et al., Supramodality of neural entrainment: Rhythmic visual stimulation causally enhances auditory working memory performance. Sci Adv, 2022. 8(8): p. eabj9782.

28. Donoghue, T., et al., Parameterizing neural power spectra into periodic and aperiodic components. Nat Neurosci, 2020. 23(12): p. 1655–1665.

29. Lu, R., et al., Aperiodic and oscillatory systems underpinning human domain-general cognition. Communications Biology, 2024. 7: p. 1643.

30. Lu, R., E. Pollitt, and A. Woolgar, Distinct and complementary mechanisms of oscillatory and aperiodic alpha activity in visuospatial attention. Imaging Neuroscience, 2025. 3.

31. Wolff, M.J., et al., Dynamic hidden states underlying working-memory-guided behavior. Nat Neurosci, 2017. 20(6): p. 864–871.

32. Barbosa, J., et al., Interplay between persistent activity and activity-silent dynamics in the prefrontal cortex underlies serial biases in working memory. Nat Neurosci, 2020. 23(8): p. 1016–1024.

33. Trubutschek, D., et al., Probing the limits of activity-silent non-conscious working memory. Proc Natl Acad Sci U S A, 2019. 116(28): p. 14358–14367.

34. Stokes, M.G., ’Activity-silent’ working memory in prefrontal cortex: a dynamic coding framework. Trends Cogn Sci, 2015. 19(7): p. 394–405.

35. Rose, N.S., et al., Reactivation of latent working memories with transcranial magnetic stimulation. Science, 2016. 354(6316): p. 1136–39.

36. Duncan, D.H., D. van Moorselaar, and J. Theeuwes, Pinging the brain to reveal the hidden attentional priority map using encephalography. Nat Commun, 2023. 14(1): p. 4749.

37. Barbosa, J., D. Lozano-Soldevilla, and A. Compte, Pinging the brain with visual impulses reveals electrically active, not activity-silent, working memories. PLoS Biol, 2021. 19(10): p. e3001436.

38. Karimi-Rouzbahani, H., et al., Temporal Variabilities Provide Additional Category-Related Information in Object Category Decoding: A Systematic Comparison of Informative EEG Features. Neural Comput, 2021. 33(11): p. 3027–3072.

39. Karimi-Rouzbahani, H. and A. Woolgar, When the Whole Is Less Than the Sum of Its Parts: Maximum Object Category Information and Behavioral Prediction in Multiscale Activation Patterns. Front Neurosci, 2022. 16: p. 825746.

40. Roy, Y., et al., Deep learning-based electroencephalography analysis: a systematic review. J Neural Eng, 2019. 16(5): p. 051001.

41. Varoquaux, G., Cross-validation failure: Small sample sizes lead to large error bars. Neuroimage, 2018. 180(Pt A): p. 68–77.

42. Jaeger, H., The “echo state” approach to analysing and training recurrent neural networks, in German national research center for information technology gmd technical report. 2001: Bonn, Germany.

43. Maass, W., T. Natschläger, and H. Markram, Real-Time Computing Without Stable States: A New Framework for Neural Computation Based on Perturbations. Neural Comput, 2002.

44. Yan, M., et al., Emerging opportunities and challenges for the future of reservoir computing. Nat Commun, 2024. 15(1): p. 2056.

45. Verstraeten, D., et al. The unified Reservoir Computing concept and its digital hardware implementations. in 2006 EPFL LATSIS Symposium. 2006. EPFL, Lausanne.

46. Suarez, L.E., et al., Connectome-based reservoir computing with the conn2res toolbox. Nat Commun, 2024. 15(1): p. 656.

47. Li, G., S. Li, and X.J. Wang, A hierarchy of time constants and reliable signal propagation in the marmoset cerebral cortex. Nat Commun, 2025. 16(1): p. 11640.

48. Dahmen, D., et al., How heterogeneity shapes dynamics and computation in the brain. Neuron, 2025.

49. Spitmaan, M., et al., Multiple timescales of neural dynamics and integration of task-relevant signals across cortex. Proc Natl Acad Sci U S A, 2020. 117(36): p. 22522–22531.

50. Bernacchia, A., et al., A reservoir of time constants for memory traces in cortical neurons. Nat Neurosci, 2011. 14(3): p. 366–72.

51. Tangermann, M., et al., Review of the BCI Competition IV. Front Neurosci, 2012. 6: p. 55.

52. Grossberg, S., Recurrent neural networks. Scholarpedia, 2013. 8: p. 1888.

53. Graves, A., Long Short-Term Memory, in Supervised Sequence Labelling with Recurrent Neural Networks, A. Graves, Editor. 2012, Springer Berlin Heidelberg: Berlin, Heidelberg. p. 37–45.

54. Ashish Vaswani, et al., Attention Is All You Need, in Advances in Neural Information Processing Systems. 2017.

55. Lawhern, V.J., et al., EEGNet: a compact convolutional neural network for EEG-based brain–computer interfaces. Journal of Neural Engineering, 2018. 15(5): p. 056013.

56. Cohen, M.X., Analyzing Neural Time Series Data: Theory and Practice. 2014, The MIT Press.

57. Jeon, Y., et al., Event-related (De)synchronization (ERD/ERS) during motor imagery tasks: Implications for brain–computer interfaces. International Journal of Industrial Ergonomics, 2011. 41(5): p. 428–436.

58. Song, Y., et al., EEG Conformer: Convolutional Transformer for EEG Decoding and Visualization. IEEE Trans Neural Syst Rehabil Eng, 2023. 31: p. 710–719.

59. Hämäläinen, M., et al., Magnetoencephalography—theory, instrumentation, and applications to noninvasive studies of the working human brain. Reviews of Modern Physics, 1993. 65(2): p. 413–497.

60. Hauk, O., M. Stenroos, and M.S. Treder, Towards an objective evaluation of EEG/MEG source estimation methods - The linear approach. Neuroimage, 2022. 255: p. 119177.

